# RES complex is associated with intron definition and required for zebrafish early embryogenesis

**DOI:** 10.1101/197939

**Authors:** Juan P. Fernandez, Miguel A. Moreno-Mateos, Andre Gohr, Shun Hang Chan, Manuel Irimia, Antonio J. Giraldez

## Abstract

Pre-mRNA splicing is a critical step of gene expression in eukaryotes. Transcriptome-wide splicing patterns are complex and primarily regulated by a diverse set of recognition elements and associated RNA-binding proteins. The retention and splicing (RES) complex is formed by three different proteins (Bud13p, Pml1p and Snu17p) and is involved in splicing in yeast. However, the importance of the RES complex for vertebrate splicing, the intronic features associated with its activity, and its role in development are unknown. In this study, we have generated loss-of-function mutants for the three components of the RES complex in zebrafish and showed that they are required during early development. The mutants showed a marked neural phenotype with increased cell death in the brain and a decrease in differentiated neurons. Transcriptomic analysis of *bud13*, *snip1* (*pml1*) and *rbmx2* (*snu17*) mutants revealed a global defect in intron splicing, with strong mis-splicing of a subset of introns. We found these RES-dependent introns were short, rich in GC and flanked by GC depleted exons, all of which are features associated with intron definition. Using these features we developed a predictive model that classifies RES dependent introns. Altogether, our study uncovers the essential role of the RES complex during vertebrate development and provides new insights into its function during splicing.

---

Splicing is critical step in eukaryotic gene expression and is an important source of transcriptomic complexity (Licatalosi and Darnell 2010). Splicing is carried out by the spliceosome, a large macromolecular complex that includes five small nuclear ribonucleoproteins (snRNPs; U1, U2, U4, U5 and U6) and hundreds of core and accessory proteins that ensure the accurate removal of introns from pre-mRNAs (Wahl et al. 2009). Canonically spliced introns are removed through two transesterification reactions during a complex process involving the recruitment and release of multiple core splicing factors (Will and Luhrmann 2011). However, many introns are recognized by different mechanisms depending on their specific features. Short introns with high GC content are believed to be spliced through an “intron definition” mechanism, in which initial U1-U2 pairing occurs across the intron. On the other hand, long introns surrounding short exons are recognized and spliced through “exon definition” mechanisms, in which the initial pairing bridges across the exon (De Conti et al. 2013). While these mechanisms are widely accepted, little is known about the specific factors associated with each process.

The pre-mRNA <u>RE</u>tention and <u>S</u>plicing (RES) complex is a spliceosomal complex conserved from yeast to human. It is organized around the U2 snRNP-associated protein Snu17p/Ist3p (RBMX2 in human), which binds to both the pre-mRNA-leakage protein 1 (Pml1p; SNIP1 in human) and bud site-selection protein 13 (Bud13p; BUD13 in human) (Gottschalk et al. 2001; Dziembowski et al. 2004; Bessonov et al. 2010). Snu17p interacts directly with the pre-mRNA and with Bud13p, and both proteins have been involved in splicing; on the other hand, Pml1p has been mainly linked to the retention of unspliced pre-mRNA in the nucleus (Gottschalk et al. 2001; Dziembowski et al. 2004; Brooks et al. 2009; Tuo et al. 2012). Furthermore, the components of the RES complex cooperatively increase the stability and the binding affinity of the complex for the pre-mRNA (Tripsianes et al. 2014; Wysoczanski et al. 2014; Wysoczanski et al. 2015; Wysoczanski and Zweckstetter 2016), highlighting the importance of cooperative folding and binding in the functional organization of the spliceosome (Tripsianes et al. 2014; Wysoczanski et al. 2014; Wysoczanski et al. 2015).

In yeast, the RES complex interacts with the 3'end of the intron in the actin pre-mRNA and is required for the first catalytic step of splicing (Gottschalk et al. 2001; Wysoczanski and Zweckstetter 2016). Additionally, microarray-based studies have shown a global effect of the RES complex on yeast splicing (Clark et al. 2002; Khanna et al. 2009). Mutations in the RES-complex genetically interact with other spliceosomal components, and several introns seem particularly sensitive to RES complex loss-of-function, often in association with weaker splice sites (Dziembowski et al. 2004; Spingola et al. 2004; Schmidlin et al. 2008; Tuo et al. 2012; Zhou et al. 2013; Wysoczanski and Zweckstetter 2016). Interestingly, disruption of the genes encoding for the three subunits of the RES complex show consistent phenotypes including slow growth, thermosensitivity (Dziembowski et al. 2004) and alteration in budding pattern (Ni and Snyder 2001).

While the RES complex was identified in yeast, its function in vertebrates, the features recognized by this complex and its role during development are unknown. Here, we found that expression of the RES complex is enriched in the CNS during early development. We generated loss-of-function mutants for the three components of the RES complex in zebrafish using an optimized CRISPR-Cas9 gene editing system (Moreno-Mateos et al. 2015). The three mutants showed severe brain defects with a significant decrease in the number of differentiated neurons and increased cell death in the brain and the spinal cord. We observed a mild retention across most introns, consistent with a global effect on splicing. However, a subset of introns was strongly affected. Importantly, these retained introns showed the hallmarks of intron definition (Amit et al. 2012; De Conti et al. 2013), as they: (i) were shorter, (ii) had a higher GC content, and (iii) were neighbored by lower GC content exons. We developed a logistic regression model with these and other genomic characteristics that allowed us to discriminate between RES-dependent and independent introns with high accuracy. Altogether, these results provide new insights into the function of the RES complex and identify the features associated with RES-dependent splicing.

## Results

### RES complex is essential for zebrafish early embryogenesis

To determine the role of the RES complex during vertebrate development, we first analyzed its expression pattern during development. *bud13*, *rbmx2* and *snip1* (Fig. 1A) are maternally expressed (Fig. 1B; Supplemental Fig. S1D), and later in development their mRNAs are strongly expressed in the central nervous system (CNS) (26 hours post fertilization, hpf), (Fig. 1B) suggesting that RES complex may be required for brain development. Next, we generated mutant zebrafish lines for *rbmx2*, *snip1* and *bud13* using an optimized CRISPR-Cas9 system (Supplemental Fig. S1A) (Moreno-Mateos et al. 2015). We identified a seven-nucleotide deletion in *bud13* (*bud13*^***Δ****7/****Δ****7*^), a sixteen-nucleotide deletion in *rbmx2* (*rbmx2*^***Δ****16/****Δ****16*^) and eleven-nucleotide deletion in *snip1* (*snip1*^***Δ****11/****Δ****11*^). These mutations are predicted to cause premature stop codons and disrupt protein function (Fig. 1C-E). Zygotic mutants for the three components of the RES complex showed a Mendelian ratio of homozygous mutant embryos. We observed strong structural brain defects and widespread cell death in the CNS at ∽30 hpf in *bud13* and ∽48 hpf in *rbmx2* and *snip1* (Fig. 1F-H; Supplemental Fig. S1B, C, E, S2B). The embryo progressively degenerates and mutants die by 4-5 dpf. Earlier depletion of *bud13* gene expression using morpholino antisense oligonucleotide targeting the AUG start site showed a more severe phenotype affecting not only the brain and CNS but also other tissues (e.g. mesoderm) (Supplemental Fig. S2A), consistent with a repression of the maternal contribution in the morphants compared to the zygotic mutants. Both, the mutant and morphant phenotypes were specific and fully rescued by injection of the cognate mRNA (*bud13*, *rbmx2 or snip1*) (Fig. 1F-H) or human mRNA (h*BUD13*) up to 6 dpf (Supplemental Fig. S2B). Interestingly, rescuing with lower amounts of h*BUD13* mRNA partially phenocopied the *rbmx2* and *snip1* mutant phenotypes at 48 hpf (Supplemental Fig. S2C). These results suggest that the onset of the zygotic phenotype in *rbmx2* and *snip1* mutants (48 hpf) is likely due to differences in the maternal mRNA contribution and/or protein stabilities for these genes. Taken together, these results demonstrate that (i) the mutant phenotypes are specific to the targeted loci, (ii) the RES complex is essential for embryonic development, and that (iii) the biochemical function of Bud13, and presumably of the RES complex, is conserved from human to zebrafish.

**Figure 1.**
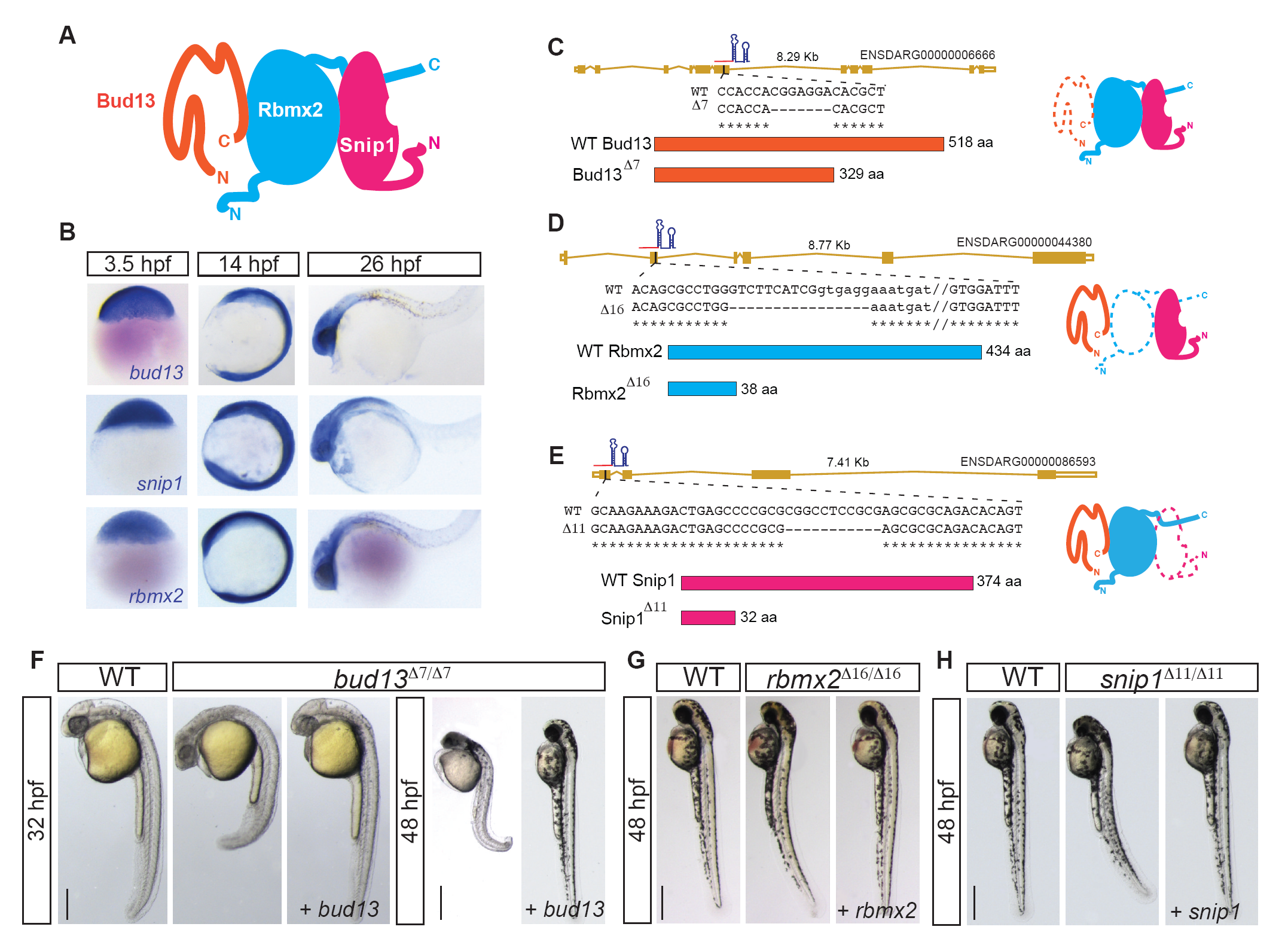
RES complex is essential for early vertebrate development. A) Schematic model of the RES complex adapted from Brooks et al. 2008. Rbmx2 (light blue) is the core subunit with an RRM-domain structure. Bud13 (orange) and Snip1 (pink) interact with Rbmx2 (light blue). B) In situ hybridization showing spatial and temporal expression of RES complex members. CE) Gene models of the mutant allele generated using CRISPR/Cas9-nanos. C) *bud13* 7 nt deletion in exon 6 generated a premature stop codon. D) *rbmx2* 16 nt deletion removed exon-intron boundary at exon 2 (exon capital letter, intron lower letter). E) *snip1* 11 nt deletion in exon 1 generated a premature stop codon. F-G) Lateral view of RES complex mutant embryos, their corresponding WT sibling and mutants injected with the cognate mRNA. F) *bud13* mutant at 32 hpf (scale bar: 0.5mm) and at 48 hpf (scale bar: 1mm). G, H) *rbmx2* and *snip1* mutant at 48 hpf respectively. WT: represent phenotypically wild type sibling from the same mutant fish line.

To determine the role of the RES-complex in splicing and gene expression, we analyzed polyA+ RNA from each mutant at the onset of the phenotype. As a control, we analyzed the transcriptome of stage-matched wild type siblings (See material and methods). We observed 621 up-regulated genes in all three mutants, that were significantly enriched for functions related to i) cell death (e.g. *p53*, *casp8* and *puma*) (Supplemental Fig. S4D), consistent with the appearance of apoptosis in the brain (Fig. 2A; Supplemental Fig. S1B), and ii) spliceosomal components, suggesting a compensatory effect upon a general splicing deficiency (See Materials and Methods for details, Supplemental Fig. S4A-C and Supplemental Table S1). In contrast, 745 genes were consistently down-regulated and were enriched for genes involved in transcriptional regulation (such as *sox19b*, *atoh7*, *pou3f1*) and nervous system development (e.g. *neurod1/4/6b*, *sox1a*) (Supplemental Fig. S4B, Supplemental Table S1).

**Figure 2.**
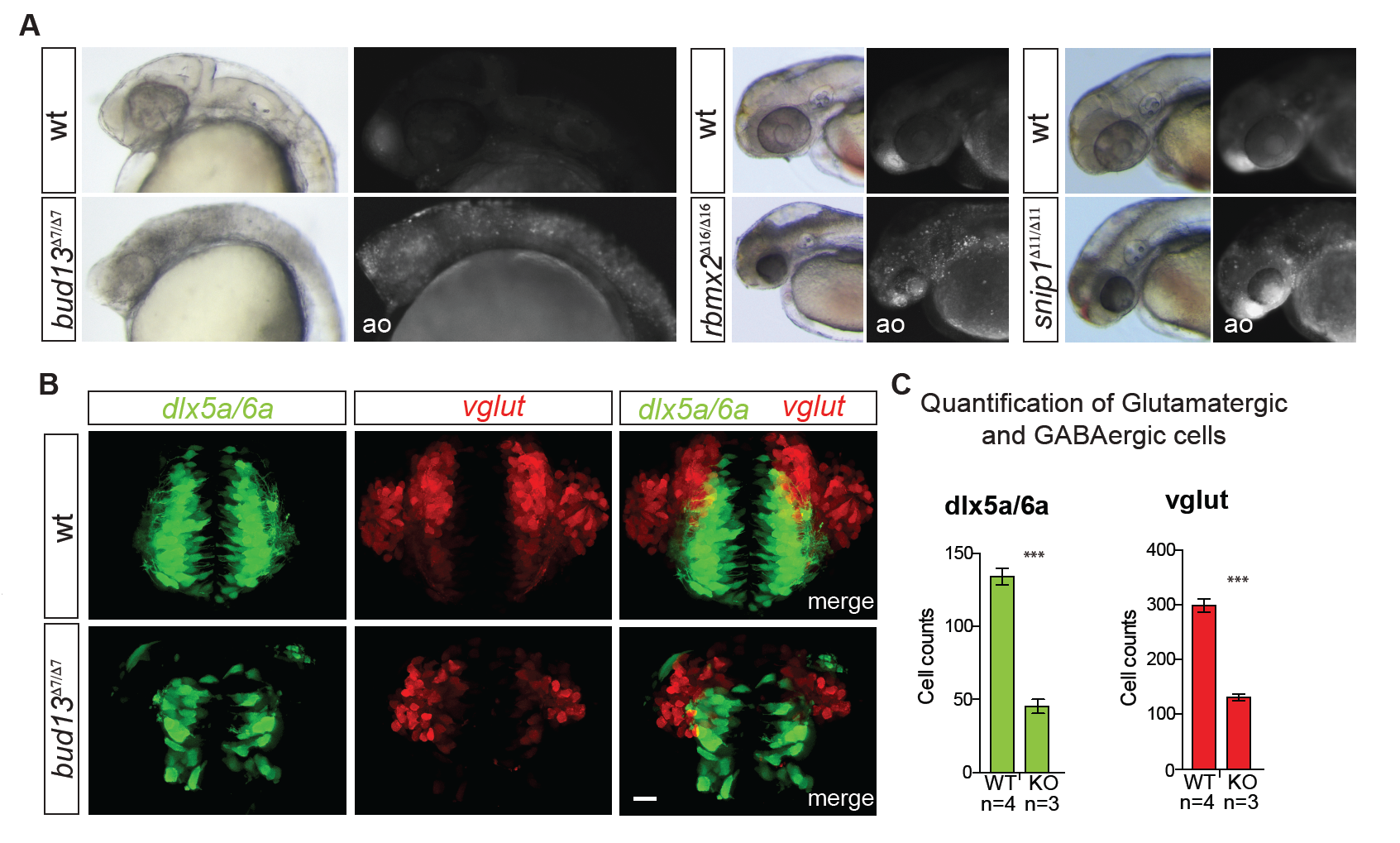
RES complex is required during zebrafish brain development. A) Acridine orange (ao) staining of zebrafish mutant embryos for *bud13* (30-32 hpf), *rbmx2* and *snip1* (48 hpf).Mutants show a marked degree of cells with nuclear uptake of ao compared to WT sibling most predominantly in the head. WT: represent phenotypically wild type sibling from the same mutant fish line. B) Maximum intensity projections of individual and merged channels (GFP and dsRed) of 3D confocal images of *bud13*^Δ7/ Δ7^ and their wild-type siblings (scale bar 20 μm) in transgenic lines that label GAB Aergic neurons and precursors (Tg [dlx6a-1.4kbdlx5a/dlx6a: GFP]) and glutamatergic neurons (Tg [vglut: DsRed]). WT: represent phenotypically wild type sibling from the same mutant fish line. C) Total number of dlx5a/6a: GFP+ cells (GABAergic neurons and precursors) and vglut:DsRed+ cells (glutamatergic neurons) in the forebrain of the *bud13*^Δ7/ Δ7^ (n = 3) and WT sibling (n = 4) were quantified. *** vglut: *P* = 2 x 10^-4^; ***dlx5a/6a: *P* = 3 x 10^-4^ (one-way ANOVA).

Importantly, *bud13*^***Δ****7/****Δ****7*^ mutants showed normal neural induction, morphogenesis and regionalization. For example, key brain areas such as the *zona limitans intrathalamica* (ZLI; dorsal *shh* in the diencephalon), the mid/hind-brain boundary (*pax2a* expression), and rhombomeres 3 and 5 (*krox20* expression) were properly specified (Supplemental Fig. S3). This suggests that the zygotic function of *bud13* is not required for initiation of neural linage patterning and specification. In contrast, we observed a reduction in the number of differentiated neurons, including both excitatory and inhibitory neuronal populations, with significant decrease in glutamatergic as well as GABAergic neurons in the forebrain in *bud13*^***Δ****7/****Δ****7*^ embryos at 32 hpf (Fig. 2B, C). These results suggest that zygotic RES activity is required for neuronal differentiation and/or survival, but not for neural induction and early brain patterning.

### RES complex mutants show widespread intron mis-splicing

To determine the global impact of the RES complex mutants on splicing, we analyzed the level of intron retention (IR) across the transcriptome. Briefly, for any given intron, the percent intron retention (PIR) is calculated as the average number of reads mapping to the 5' and 3' Exon-Intron (E-I) junctions over the average number of reads mapping E-I junctions plus any Exon-Exon (E-E) junction that supports removal of that given intron (Fig. 3C) (Braunschweig et al. 2014). We found that 74-79% of introns showed increased retention (ΔPIR > 0) in the individual mutants compared to wild type siblings (out of 72,926 introns, with sufficient coverage, see Methods for details). In contrast, a significantly smaller set of introns (7-9%) show decreased retention in the mutants (Fig. 3A, B, *P* < 2.2e-16, Fisher exact test). Interestingly, the three mutants shared 5,339 (∽35%) introns with a medium-high level of retention (ΔPIR > 5) (Fig 3D), consistent with a common function in the RES complex. This is likely an underestimate because each mutant was analyzed at the onset of the mutant phenotype, which is different across these mutants likely due to different level of maternal recue or protein stability. despite the fact that the onset of the phenotype was different. We observed mild widespread intron retention, yet ∽3.5% of introns showed strong increased retention across each mutant (ΔPIR >15, Fig 3B), suggesting that a subset of introns have a stronger dependence for RES function *in vivo*. This effect was validated by RT-PCR for a subset of candidates for each mutant (Fig 3F, Supplemental Fig. S5). Genes with strongly retained introns (ΔPIR > 15) in at least two of the three mutants were enriched in transcriptional regulation and DNA binding (Supplemental Fig. S6), including known regulators of vertebrate development and neuronal differentiation (e.g. *irx1a*, *smn1*, *enc1, smad4*, *tbx2a*, and *nkx6.1/6.2*; Supplemental Table S2). Taken together, our analyses indicate that the RES complex has a global effect on splicing, and is strongly required for a subset of introns in genes involved in transcriptional regulation and neural development.

**Figure 3.**
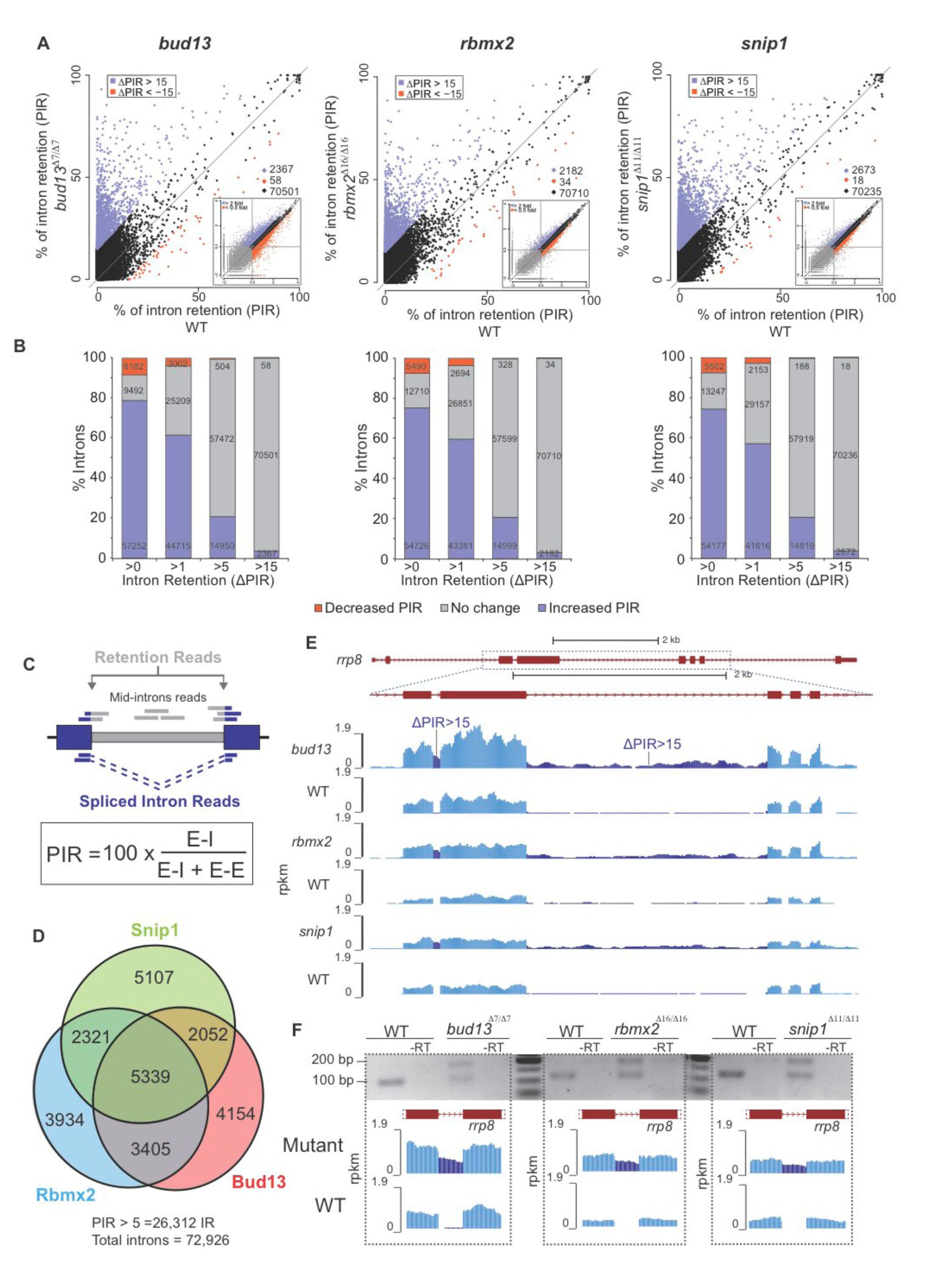
RES complex mutants show mild widespread intron mis-splicing. A) Biplot illustrating percent intron retention (PIR) in each of the RES mutants and the corresponding phenotypically wild type (WT) siblings. Blue and red dots correspond to introns with higher inclusion in the mutant and WT, respectively, using a cutoff of ΔPIR > 15. Insets show exon gene expression levels in the same conditions (see Supplemental Fig. S4 for details). B) Stacked barplot showing the percentage of introns affected by *bud13*, *rbmx2* or *snip1* mutation using different ΔPIR cutoffs. C) Scheme showing how PIR was measured (adapted from Braunschweig et al. 2014; see Methods for details). D) Euler diagram showing the number of retained introns (ΔPIR >5) in the three mutants and the inter-mutant overlaps. E) RNA-seq read density across the *rrp8* gene in *bud13*, *rbmx2* and *snip1* mutants and their corresponding phenotypically WT siblings. Intronic signal increases in RES mutants (ΔPIR>15) (dark blue) (dotted square box). F) RT-PCR assays validate the increased retention of an *rrp8* intron (from panel E) in *bud13*, *rbmx2* and *snip1* mutants compared to the corresponding phenotypically WT siblings.

### Distinguishing features and classes of retained introns in the RES complex

To identify features that are primarily associated with RES complex function, we first defined a set of introns that were confidently dependent on RES for proper splicing (1,409 “RESdep” introns with ΔPIR>15, ≥1.5-fold net increase in intron reads; see Methods) (Fig. 3C). As a control set (Ctr), we defined 5,574 introns with ΔPIR<0.5 in all three mutants, and evaluated the enrichment of 44 features, many of which have been previously associated with intron retention (Braunschweig et al. 2014) (Supplemental Table S3). Consistent with the genome-wide patterns (Fig. 4), strongly retained introns were enriched for last introns (Fig. 4A, B, Supplemental Fig. S7A) and introns that do not trigger NMD when mis-spliced (Fig. 4A, C, Supplemental Fig. S7B), suggesting that their accumulation is in part likely due to reduced degradation of the unspliced transcript isoform. However, a significant fraction of highly retained introns was predicted to elicit NMD upon inclusion. We thus hypothesized that this subset of NMD-triggering introns contains specific features that would maximally associate with RES-dependent mechanisms. Based on this, we also separately analyzed introns that were predicted to trigger NMD (574) and those that were not (569) (see Methods).

**Figure 4.**
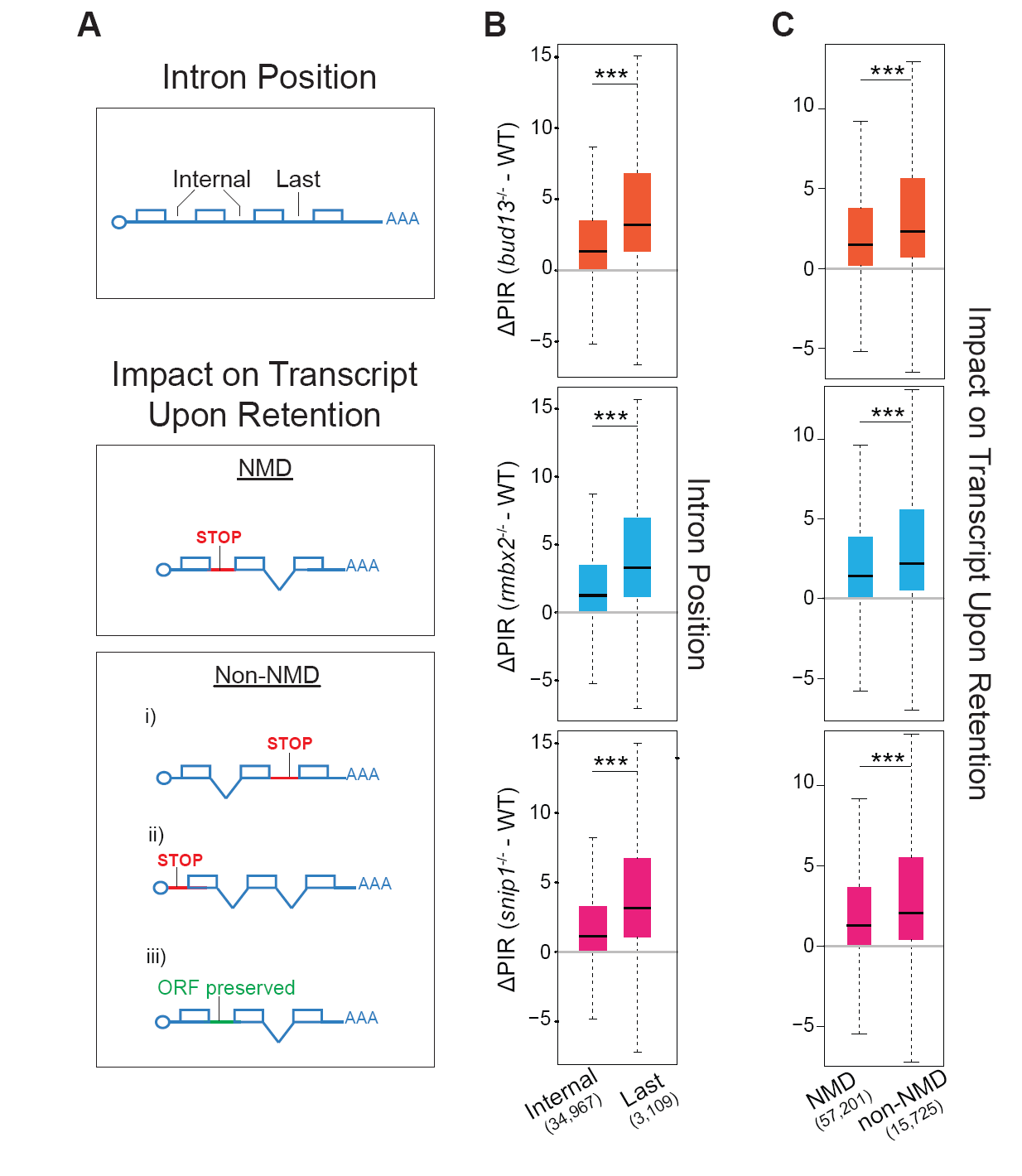
Genome-wide analysis of intron retention in RES complex mutants A) Schematic representation of intron features analyzed in B and C. Top: Intron position. “Internal” introns correspond to all introns excluding the first two and last three introns. Bottom: NMD vs non-NMD triggering introns. Introns predicted to cause NMD upon retention introduce a premature termination codon (PTC) further than 50 nucleotides upstream of an exon-exon junction. Introns predicted not to cause NMD (noNMD) may correspond to: (i) last introns, (ii) introns in UTRs or non-coding genes, or (iii) introns that preserved the ORF upon retention (multiple of three nucleotides with no in-frame stop codons). B) Box plots showing the ΔPIR of last and internal introns in the three different mutant of the RES complex. Only genes with more than 10 introns were considered for the analysis. *bud13* *** *P* = 4. 3 7 x 1 0^-201^; *rbmx2* * * * *P* =2.98x10^-202^; *snip1* *** *P* =4.76x10^-195^ (Wilcoxon rank sum test). C) Box plots showing the ΔPIR of introns predicted to trigger nonsense mediated decay (NMD) upon retention and those predicted not to trigger NMD (no-NMD) in the three different mutant of the RES complex. *bud13* *** *P* = 5.83x10^-208^; *rbmx2* *** *P* =4.94x10^-184^; *snip1* *** *P* =1.74x10^-183^ (Wilcoxon rank sum test).

RES-dependent introns (i) were significantly shorter than the control set (Fig. 5A, median of 281 nt versus 749 nt in the control, *P*<0.001, Mann-Whitney-U test), (ii) had elevated GC content (Fig 5B, *P*<0.001 Mann-Whitney-U test), (iii) were flanked by exons with lower GC content (Fig. 5C, D; Supplemental Fig. S8C, D), and (iv) had weaker branch point (BP) consensus sequences (Corvelo et al. 2010; Corioni et al. 2011) (Fig. 5G; Supplemental Fig. S8). Remarkably, at the genome-wide level, short introns and introns with high GC content also showed a higher retention across all three mutants compared to wild type siblings (Supplemental Fig. S9). NMD-triggering introns further showed weaker core splicing signals, including acceptor (3') and donor (5') splice sites and BP consensus sequences than any other intron set (Fig. 5E-G), providing an explanation for their increased sensitivity upon RES complex disruption despite being subject to NMD.

**Figure 5.**
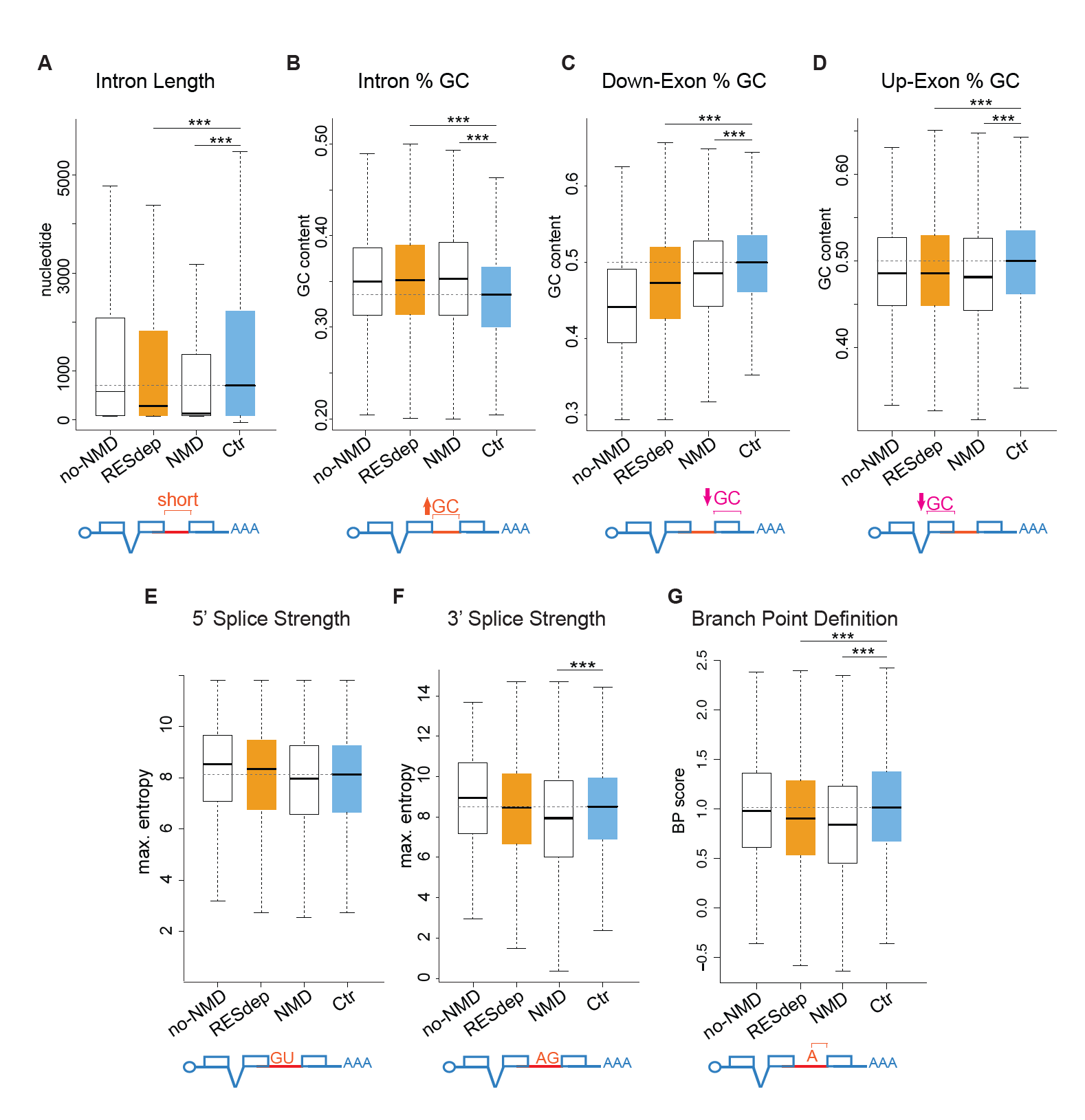
Molecular features that determine intron retention in the RES complex mutants are associated with splicing through intron definition A-G) Box plots showing the median (black solid line) and the distribution of values for multiple intron-exon features of several groups of introns of interest. “RESdep”, all highly retained introns (ΔPIR > 15) in at least two out the three mutants in the RES complex; “no-NMD”, subset of “RESdep” retained introns that are predicted not to trigger NMD; “NMD”, subset of “RESdep” retained introns predicted to trigger NMD; and “Ctr”, control set of introns with a ΔPIR cutoff < 0.5 in the three RES complex mutants (see Materials and Methods for details). (*** *P* ≤ 0.001; Mann-Whitney-U test). Branch point (BP) definition: BP Score of best-predicted BP. Scored base on Corvelo et al. 2010 (See Materials and Methods for details).

Next, we assessed how genomic and transcript features define the dependence on the RES-complex. We applied a logistic regression model using 30 features as predictor variables (Supplemental Table S3), and defined a response variable classifying each intron as RES-dependent or non-dependent. We developed a model using 90% of the RESdep and a size-matched subset of control introns as training set, and validated the model on the remaining 10% of the data (see Methods). We found a high performance in the classification of introns, with an average Area Under the ROC curve (AUC) of 0.821 (Fig. 6A). This model was able to classify the impact of mutating each individual RES component with an AUC of ≥0.76. Subsequently, we analyzed the individual contribution of each feature to the model and their potential to reduce the null deviance (see Methods). Consistent with results in Fig. 5, this analysis identified four important features i) the ratio between exon and intron length, ii) the ratio between exon and intron GC content, iii) gene expression levels and iv) the position of the intron within the transcript (last intron effect) (Fig. 6B and Supplemental Fig. S10). Altogether, our logistic regression analysis can identify introns dependent on the RES complex based on specific features within the genomic locus and the transcript.

**Figure 6.**
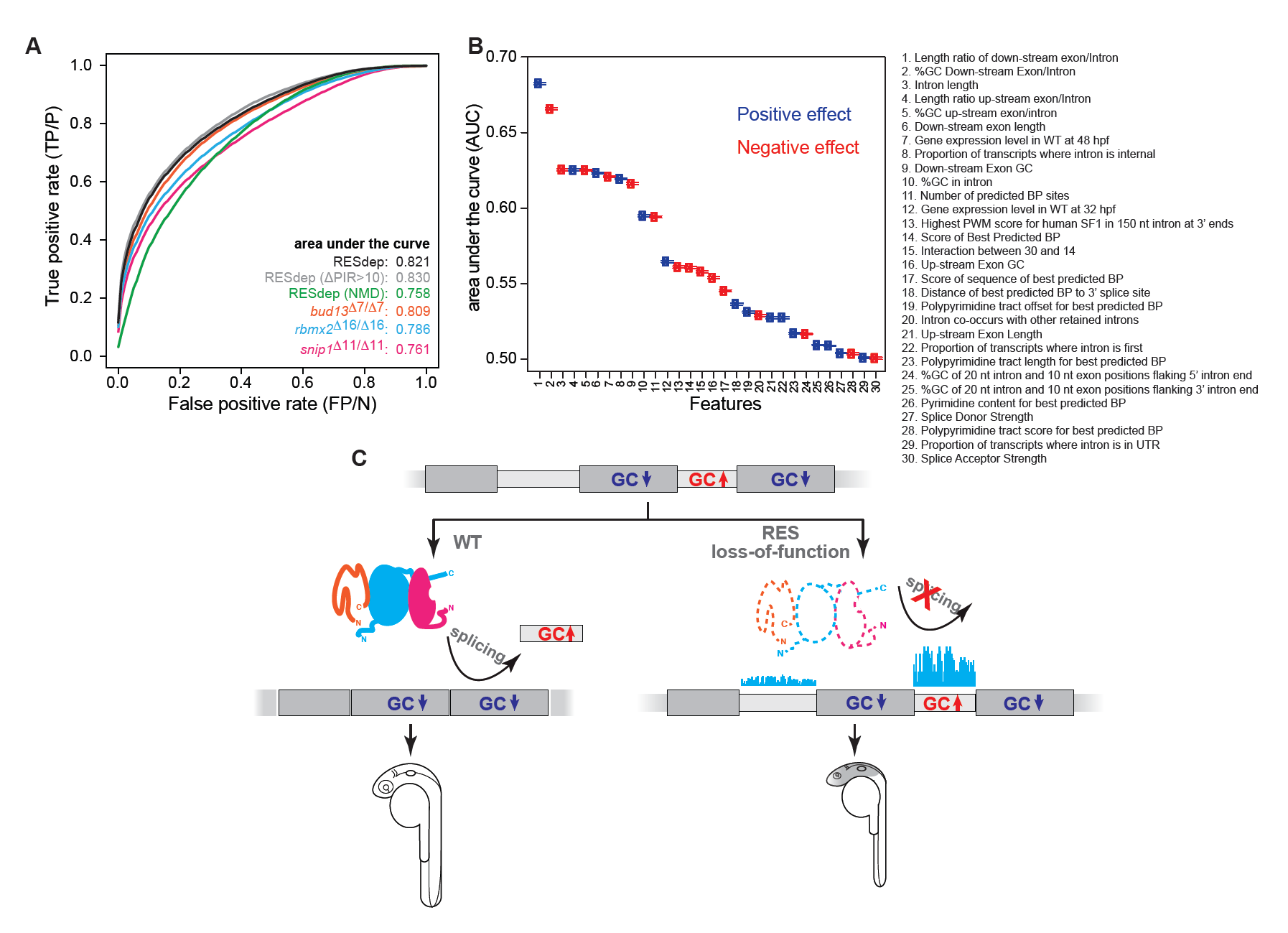
Logistic regression model can accurately classify RES-dependent and non-dependent introns. A) Classification performance of logistic regression models for different data sets of differentially retained vs. Ctr introns. ROC curves are averaged over 10,000 repeated holdout experiments where models have been trained with randomly sampled subsets of 90% (1,268) of the RESdep introns versus 1,268 Ctr introns with 30 features (Supplemental Table S3) and Lasso feature selection. Classification performance was estimated using the remaining 10% (141) RESdep introns and 141 randomly sampled Ctr introns. Having held fixed parameters, the same model was used to estimate classification performance with randomly sampled 141 introns from the other RES-dependent data sets, namely: (i) “RESdep (ΔPIR >10)” introns from the “RESdep” set with ΔPIR > 10 in all three mutants (871 introns); (ii) “RESdep (NMD)”, introns from the “RESdep” set predicted to trigger NMD when retained (574 introns); (iii) “*bud13*^*Δ7/Δ7*^”, introns with ΔPIR>15 upon *bud13* mutation at 32 hpf (2,363 introns); (iv) “*rbmx2*^*Δ16/Δ16*^”, introns with ΔPIR>15 upon *rbmx2* mutation at 48 hpf (2,186 introns); and (v) “*snip1*^*Δ11/Δ11*^”, introns with ΔPIR>15 upon *snip1* mutation at 48 hpf (2,675 introns). 95% confidence interval of reported average AUCs corresponds to AUC ± 0.001. B) Capacity of each feature to discriminate between “RESdep” and “Ctr” introns, measured as AUC (average area under ROC curve) when used as the only feature in a one-feature logistic regression model. C) Schematic integrative model of the RES complex function in splicing. The loss-function of the RES complex induces a global weak defect on splicing, but a strong retention of a subset of introns with particular features. These introns are short, have a higher GC content and flanked by GC depleted exons, features associated to intron definition splicing mechanism. This splicing defect leads to a brain related phenotype.

## Discussion

Splicing regulatory information is encoded by multiple sequence features, from the core signals (splice donor and acceptor and branch point) to other, less understood, sequence elements (Papasaikas and Valcarcel 2016; Savisaar and Hurst 2017; Vaz-Drago et al. 2017). Our results identify intronic features that are associated with RES-dependent splicing across the transcriptome. These features can be used to discriminate a large fraction of RES-dependent from independent introns. Although loss-of-function for RES complex components caused mild intron retention across the transcriptome, we observed a subset of introns that were strongly accumulated across mutants of the RES complex. Recent *in vitro* studies on the single intron of the actin gene in yeast showed that the RES complex binds at the 3’ of the intron, between the BP and the acceptor site (Wysoczanski et al., 2014). We observed that introns that more strongly depend on the RES complex show weaker BP consensus sequences. Furthermore, this subset of introns were shorter and had higher GC content, an association that is particularly striking in zebrafish, since short introns normally have lower GC content (Amit et al., 2012). RES-dependent introns are flanked by exons with a lower GC content than RES-independent introns (Fig. 5; Supplemental Fig. S9C, D). Therefore, these observations are consistent with a model whereby RES-dependent introns are mainly spliced through intron definition (Amit et al., 2012; De Conti et al., 2013). This association is surprising, since biochemical evidence suggest that the RES complex joins the spliceosome after recognition of the splice sites (Wysoczanski et al., 2014), and that the RES complex is not needed for spliceosome assembly *in vitro* but for U1 and U4 snRNP dissociation before the first catalytic step (Gottschalk et al. 2001). One possible explanation for this apparent discrepancy is that RES complex components play a role in early splice site recognition *in vivo* and therefore that the biochemical functions reported in a limited set of RNAs reflect limitations of *in vitro* splicing reactions. Alternatively, the RES complex may not be involved in early splice site recognition, but could be a limiting factor for splice site pairing or other steps in spliceosome assembly progression *in vivo*, particularly for introns defined by intron definition, highlighting differences in molecular pathways for intron-and exon-defined splicing. These concepts are in line with increasing evidence that splice site selection can be modulated at late stages of spliceosome assembly or even catalysis (Clark et al. 2002; Park and Graveley 2005; Pleiss et al. 2007; Hoskins and Moore 2012; Chen and Moore 2014; Papasaikas et al. 2015; Tejedor et al. 2015; Papasaikas and Valcarcel 2016). Finally, some RES-dependent introns also have weaker donor and acceptor splice site consensus sequences, and thus are expected to be more sensitive to defects on the splicing machinery. This is consistent with previous studies in yeast, which found that weaker 5’ splice sites increased susceptibility to RES loss-of-function (Dziembowski et al., 2004).

An unexpected observation from genome-wide analyses of core splicing factor loss-of-function experiments is that each factor seems to differentially affect a specific subset of introns and exons (Saltzman et al. 2011; Papasaikas et al. 2015). This suggests that splicing of each intron in the genome is limited by specific core factors, depending on its combination of sequence features, as we observed for RES-dependent introns. As such, disruption of core splicing factors is predicted to produce unique phenotypes dictated by its expression, and the expression and function of genes that contain the subsets of introns sensitive to that factor. Consistent with this hypothesis, RES complex is required during brain development and neuronal survival, and mis-regulated introns are found in genes with well-known functions in neurodevelopment (e.g. *irx1a*, *smn1*, *enc1, smad4*, *tbx2a*, and *nkx6.1/6.2*). Specifically, zygotic mutants in *bud13*, *snip1* or *rbmx2* show microcephaly and decreased populations of GABAergic and glutamatergic neurons, despite normal specification and regionalization of the CNS (Fig. 1, Fig. 2 and Supplemental Fig. S3). This phenotype is different from those described for a few other spliceosomal-related mutants in zebrafish (An and Henion 2012; English et al. 2012; Keightley et al. 2013; De La Garza et al. 2016). For instance, while *sf3b1* is required for early neural crest development (An and Henion 2012), loss of another core component of the spliceosome, *prpf8*, results in massive neuronal cell death and impaired myeloid differentiation (Keightley et al. 2013). These differences might be caused by the different half-life of the maternal proteins in the zygotic mutants. Alternatively, different components of the splicing machinery might be essential in a cell – type/tissue specific manner during early development. This may also explain why mutations in specific spliceosomal components cause human diseases with diverse phenotypes, such as Taybi-Linder syndrome, microcephalic osteodysplastic primordial dwarfism type I and retinitis pigmentosa (Towns et al. 2010; Edery et al. 2011; He et al. 2011). Interestingly, a mutation in human *SNIP1* (p.Glu366Gly) has been associated with epilepsy and skull dysplasia (Puffenberger et al. 2012). Our data shows that human *BUD13* can rescue loss of *bud13* function in zebrafish, and future studies will be needed to determine whether Bud13 has a conserved function during brain development in humans (Supplemental Fig. S2B, C).

In summary, we have shown that RES complex disruption in zebrafish hinders splicing, but is not essential for the removal of most introns, indicating that such introns can be efficiently defined and spliced through RES-independent mechanisms. However, we found that a subset of introns is particularly affected by RES complex removal and that those introns display the major hallmarks of splicing through intron definition mechanisms. From a functional perspective, RES-dependent introns are in genes enriched for transcription factors and neurodevelopmental regulatory functions, thus resulting in brain developmental defects in loss-of-function zygotic mutants. Future studies will be needed to understand how spliceosomal mutations disrupt splicing of different genes by affecting specific limiting steps in pre-mRNA splicing resulting in diverse disease phenotypes.

## Methods

### Zebrafish maintenance, mating and embryos image acquisition

Fish lines were maintained in accordance with research guidelines of the International Association for Assessment and Accreditation of Laboratory Animal Care, under a protocol approved by the Yale University Institutional Animal Care and Use Committee (IACUC). Wild-type zebrafish embryos were obtained through natural mating of TU-AB and TLF strains of mixed ages (5 – 17 months). Selection of mating pairs was random from a pool of 48 males and 48 females allocated for a given day of the month. *bud13*^*Δ7/Δ7*^, *rbmx2*^*Δ16/Δ16*^, *snip1*^*Δ11/Δ11*^ were obtained through natural mating of heterozygous *bud13*^*+/Δ7*^, *rbmx2*^*+/Δ16*^, *snip1*^*+/Δ11*^ mutants, respectively (see below gene editing using CRISPRCas9). Tg(*dlx6a-1.4kbdlx5a/dlx6a:GFP*) lines (Zerucha et al. 2000; Ghanem et al. 2003) were obtained from the laboratory of Marc Ekker and Tg(*vglut:DsRed*) (Kinkhabwala et al. 2011) from the laboratory of Joseph Fetcho.

Embryos were analyzed using a Zeiss Axioimager M1 and Discovery microscopes and photographed with a Zeiss Axiocam digital camera. Images were processed with Zeiss AxioVision 3.0.6.

### Confocal microscopy imaging and Data Processing

Whole-mount embryos were imaged in vivo by confocal microscopy (Leica TCS SP8 systems, Yale Center for Cellular and Molecular Imaging.) Mutant embryos and their wild-type siblings were scored at 30-32 hours post fertilization stage. Embryos were anesthetized using Tricaine and mounted in 0.6% agarose at an orientation where the frontal view of the brain was imaged. Embryos were imaged at 40x (1.3 Oil) using Z-stacks ranging from 70.41-96.02 μm, with a z stepping size of 0.4 μm. Z-stacks started at the first appearance of the GABAergic cells (GFP-labeled) and ended where GABAergic cells (GFP-labeled) could no longer be visualized. Each xy plane spanned 227.94 μm with a pixel size of 0.075 μm. Maximum intensity projections were shown for all confocal images, which were processed using Fiji (Schindelin et al. 2012), Imaris (Bitplane) and Huygens deconvolution software (Scientific Volume Imaging). Figures were assembled using Illustrator (CC, Adobe). To quantify the number of glutamatergic (labeled in DsRed) and GABAergic cells (labeled in GFP) in the *bud13*^Δ7/Δ7^ mutant and their wild-type siblings respectively, two blinded raters segmented the raw z-stack images using ImarisCell module (Bitplane) and computationally counted the segmented cells in each channel (GFP and DsRed). Identical thresholds and parameters were applied to all samples for segmentation processing. Since the quantification performed by both independent raters yield consistent fold change in the respective cell counts between the *bud13*^Δ7/Δ7^ mutant and their wild-type siblings. Only one set of the analyzed results was displayed. Statistical analyses were conducted using Prism 6 (Graphpad).

### Acridine Orange Staining

To visualize apoptotic cells, vital dye acridine orange (Sigma) was used in live and dechorionated embryos. Embryos were incubated 2 minutes in PBS pH 7.1 with 2 ug/ml of acridine orange in the dark. After 3 brief washes in PBS, the embryos were placed in plates with 1% agarose and viewed with fluorescence microscopy, using the FITC filter set 1 (Brand et al. 1996).

### Plasmids constructs

Zebrafish *bud13*, *rbmx2* and *snip1* ORFs were PCR amplified (Supplemental Table S4) using cDNA from 2 and 6 hpf zebrafish embryos and cloned in pSP64T (*bud13*) in pT3TS (Jao et al. 2013) (*rbmx2* and *snip1*). *bud13* PCR product was cut using *NotI* and *EcoRI* and and ligated into pSP64T. *rbmx2* and *snip1 PCR* products were cut with *NcoI* and *SacII* and ligated into pT3TS. To optimize Kozak sequence, the forward oligonucleotide used for *rbmx2* ORF introduced extra aminoacid in the 2nd position (GGC). Human *bud13* ORF was cloned in pSCDest (Villefranc et al. 2007) using gateway^®^ gene cloning system (Thermo Fisher Scientific). Final constructs were confirmed by sequencing. To generate mRNAs, the template DNA was linearized using *XbaI* (pT3TS), *BamHI* (pSP64T) or *KpnI* (pSCDest) and capped mRNA was synthetized using the mMessage mMachine^®^ T3 (pT3TS), or SP6 (pSP64T and pSCDest) kit (Ambion), respectively and in accordance with the manufacturer’s instructions. *In vitro* transcribed mRNAs were DNAse treated and purified using the RNeasy Mini Kit (Qiagen). All mRNA rescued the mutant phenotypes when 50-100 pg were injected in one cell stage embryo.

### Morpholino injections and gene editing using CRISPR-Cas9

A morpholino targeting *bud13* mRNA start codon was obtained from Gene Tools and re-suspended in nuclease-free water. 1 nl of morpholino solution (0.6mM) was injected into wild-type dechorionated embryos at the one-cell stage.

CRISPR/Cas9-mediated gene editing was performed as described previously (Vejnar et al. 2016). Briefly, 3 different sgRNAs (20 pg each) targeting *bud13* gene (Supplemental Table S4) were co-injected together with 100 pg of mRNA coding for zebrafish codon optimized Cas9-nanos in one-cell stage embryos (Supplemental Fig. 1SA). Cas9-nanos concentrates gene editing in germ cells and increases the viability of injected embryos (Moreno-Mateos et al. 2015). F0 founders were mosaic and they were backcrossed with wild-type fish and then F1 fish were genotyped using their corresponding oligos per target site (Supplemental Table S4). Heterozygous adult fish *bud13*^*+*/Δ7^ (Fig. 1) were selected to generate *bud13*^Δ7/Δ7^ mutants. Similar approach was followed to generate *rbmx2* and *snip1* mutants but injecting 2 sgRNAs (Supplemental Table S4 and Fig. 1).

### *In situ* hybridization

*krox20*, *shh* and *pax2a* in situ probes were previously described (Krauss et al. 1992; Krauss et al. 1993; Oxtoby and Jowett 1993). Antisense digoxigenin (DIG) RNA probes were generated by in vitro transcription in 20 μl reactions consisting of 100 ng purified PCR product (8 μl), 2 μl DIG RNA labelling mix (Roche), 2 μl × 10 transcription buffer (Roche), and 2 μl T7/T3 RNA polymerase (Roche) in RNase-free water and purified using a Qiagen RNEasy kit. In situ protocol was followed as detailed previously (Giraldez et al. 2005). To reduce variability, wild-type sibling and *bud13*^Δ7/Δ7^ embryos were combined in the same tube during in situ hybridization and recognized based on their phenotype. Before photo documentation, embryos were cleared using a 2:1 benzyl benzoate:benzyl alcohol solution. Images were obtained using a Zeiss stereo Discovery V12.

### RNAseq library, Reverse transcription PCR (RT – PCR) and qPCR

Total RNA from 32 hpf *bud13*^Δ7/Δ7^, 48 hpf *rbmx2*^Δ16/Δ16^, 48 hpf *snip1*^Δ11/Δ11^ embryos and their corresponding siblings was extracted using Trizol (ten embryos per condition). Strand-specific TruSeq Illumina RNA sequencing libraries were constructed by the Yale Center for Genome Analysis. Samples were multiplexed and sequenced on Illumina HiSeq 2000/2500 machines to produce 76-nt paired-end reads.

RNA used for intron retention validation experiments was treated with TURBO DNase (Ambion) for 30 min at 37 °C and extracted using phenol chloroform. Then, Polyadenylated RNAs were purified using Oligo d(T)25 Magnetic Beads (NEB) following manufacter recommendations. cDNA was generated by reverse transcription with random hexamers using SuperscriptIII (Invitrogen). RT – PCR reactions were carried out at an annealing temperature of 59 °C for 35 cycles. Primers are listed in the Supplemental Table S4.

For the qPCR experiment, total RNA was extracted as described above. GFP and dsRED mRNAs were used as spike-in RNA controls and 1 μg of total RNA was used to generate cDNA. 5 μl from a 1/50 dilution of the cDNA reaction was used to determine the levels of p53 in a 20 μl reaction containing 1 μl of each oligo forward and reverse (10 μM) (Supplemental Table S4), using Power SYBR Green PCR Master Mix Kit (Applied Biosystems) and a ViiA 7 instrument (Applied Biosystems). PCR cycling profile consisted of incubation at 50 °C for 2 min, followed by a denaturing step at 95 °C for 10 min and 40 cycles at 95 °C for 15 s and 60 °C for 1 min. Primers are listed in the Supplemental Table S4.

### Genotyping

Zebrafish embryos or a small amount of tissue from the end of the tail were used to extract DNA (Meeker et al. 2007). Briefly, embryos or fin clipped were incubated in 80 μl of NaOH 100mM at 95 °C for 15 min producing a crude DNA extract, which was neutralized by the addition of 40 μl of 1 M Tris-HCl, pH 7.4 (Sigma-Aldrich). 1 μl of this DNA extraction was used as a template for PCR reactions using the primers described in Supplemental Table S4.

### Gene expression analyses

Gene expression levels for each condition were calculated from RNA-seq data using the cRPKM metric (corrected-for-mappability <u>R</u>eads <u>P</u>er <u>K</u>ilobasepair of uniquely mappable positions per <u>M</u>illion mapped reads (Labbe et al. 2012). For this, a reference transcript per gene was selected from the Ensembl version 80 annotation for *Danio rerio* using BioMart (25,935 genes in total, Supplemental Table S5) and uniquely mappable positions for each transcript were calculated as previously described (Labbe et al. 2012). Quantile normalization of cRPKM values was done with ‘normalizeBetweenArrays’ within the ‘limma’ package. To identify differentially expressed genes, we first filtered out genes that did not have cRPKM > 2 in all sibling control or all mutant samples and genes whose quantification was not supported by at least 50 read counts in at least 1 sample. Next, differentially expressed genes were defined as those that showed a fold change in expression of at least 1.5 in all 3 mutants and a fold change of at least 2 in 2 out of the 3 control vs. mutant individual comparisons (*bud13*, *rbmx2* and *snip1*). Gene Ontology analysis was performed with the online tool DAVID (https://david.ncifcrf.gov/. Version 6.8) using as background all genes that passed the initial filters (minimum expression and read count).

### Genome-wide analysis of intron retention

Annotated introns for each reference zebrafish transcript in Ensembl version 80 were extracted and those that overlapped with other genes were removed yielding a total of 182,017 valid introns. To calculate the percent of intron retention (PIR) for each intron in a given RNAseq sample, we used our previously described pipeline (Braunschweig et al. 2014) with the following modification: to calculate intron removal, all exon-exon junctions supporting the splicing of the intron were used and not only those formed between the two neighboring exons. This was done to avoid false positives in the case of introns associated to cassette exons or other alternative splicing events. For all analysis, only introns with sufficient read coverage across the six samples were considered (at least 15 reads supporting the inclusion of one splice site and 10 of the other, or a total of 15 reads supporting splicing of the intron).

To define the confident set of highly affected introns, potential false positives were filter out by comparing the density of the mapped reads in the introns bodies in the mutant vs the control. For this purpose, we extracted all intronic sequences and calculated the number of uniquely mappable positions per intron following a similar strategy to that used to calculate cRPKMs (Labbe et al. 2012) (see above). Specifically, every 50-nucleotide (nt) segment in 1-nucleotide sliding intronic windows was mapped to a library of full-length intronic sequences plus the whole genome, using bowtie with –m 2 –v 2 parameters (every intronic segment must map at least twice, to its own individual intron sequence and to the corresponding position in the whole genome). Segments that mapped more than twice were considered as multi-mappable positions, whereas those that did not map (e.g. due to undetermined (N) nucleotides in the assembly) were considered as non-mappable. The number of uniquely mappable positions of an intron is defined as the total number of segments minus multi-and non-mappable positions. Next, each RNA-seq sample was mapped to the same library of intronic plus full genomic sequences using –m 2 –v 2 to obtain the unique intronic reads counts. However, to minimize potential artifacts derived from the heterogeneity of the intronic sequences (e.g. high number of reads mapping to a transposable element or an expressed nested gene), if a given intronic position showed a read count more than five times higher than the median read count of the whole intron, then the read count of this position was set to 5 × median (if the median read count was 0, then the maximum read count for any given position was set to 5). Finally, ciRPKM scores were calculated (correctedfor-mappability intronic Reads Per Kilobasepair of uniquely mappable positions per Million mapped reads) for each intron and condition by dividing this number of counts by the number of uniquely mappable positions in that intron.

With this information, the set of confidently retained introns upon RES complex disruption (“RESdep” introns) were defined as those introns with ΔPIR > 15 and at least a 1.5-fold net increase in read density in the intron body calculated as:

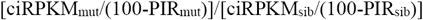

in at least 2 out of 3 mutants (1,413 introns in total). As a control, we also define a set of confidently non-retained introns as those with a ΔPIR < 0.5 in the 3 mutants (“Ctr” set; 5,577 introns).

To analyze the number of retained introns in the 3 mutants and the inter-mutant overlaps Euler APE-3.0.0 software (Micallef and Rodgers 2014) was utilized.

### Analysis of intron-associated features

To investigate the impact of non-sense mediated decay (NMD) on global intron retention upon RES mutation, all introns were separated as last or non-last introns (of the reference transcript) and between those predicted to trigger and not to trigger NMD. An intron was predicted to trigger NMD if its retention generated an in-frame stop codon that is located further than 50 nts upstream of an exon-exon junction (Hsu et al. 2017). By definition, last introns cannot trigger NMD. ΔPIR values were plotted as boxplots for each category, and two-sided Wilcoxon Sum Rank tests were used to evaluate statistical differences between the distributions.

To identify features discriminating introns highly retained upon RES depletion from un-retained/unaffected introns, we compared the sets of confidently introns (“RESdep” in Fig. 5.) with control introns (“Ctr” in Fig. 5.). Moreover, as introns that are predicted not to trigger NMD are expected to be more often accumulated unspecifically, two subsets for “RESdep” introns were generated: (i) those introns in genes with more than five introns, are not the last three introns of the gene, and that are predicted to trigger NMD (“NMD”, 577 introns); and (ii) predicted not to trigger NMD or cause a frame shift upon inclusion, unless they are the last intron of the gene (“no-NMD”, 569 introns). For these different sets of introns, 44 features were extracted (Supplemental Table S3), including intron and exon length and GC content, strength of 3' and 5' splice sites, branch point (BP) related features, and transcript length, using custom scripts in combination with the following two external tools: MaxEntScan scripts for determining the strength of 3' and 5' splice sites (Yeo and Burge 2004); and SVM-BPfinder software for determining BP related features (BP strength, distance from BP to 3' splice site, and pyrimidine track length) (Corvelo et al. 2010). For the latter analysis, the 150 nts upstream of the 3' splice sites were extracted and these sequences were used as input for SVM-BPfinder. Furthermore, we recorded the highest log-score of the SF1 position weight matrix binding model across these 150-nt intronic sequences (Corioni et al. 2011).

### Logistic regression model and evaluation of feature discrimination capacity

We applied logistic regression models to the discrimination between differentially retained introns (retained) and non-differentially retained introns (control) upon RES mutations. We focused on the set of confidently retained introns (“RESdep”, 1,409 introns) versus control introns with an absolute ΔPIR < 0.5 in the three mutants (“Ctr”, 5,565; we removed 9 introns for which we could not determine all features). The binary response variable of the logistic regression models indicates for each intron if it belongs to the retrained or control group. As predictors we used 30 quantitative and qualitative features (Supplemental Table S3), including intronic and exonic characteristics, position along the transcript and gene expression in wild type conditions, among others. The binomial logistic regression models were learned using Lasso variable selection (Friedman et al. 2010; Tibshirani et al. 2012; Simon et al. 2013) available in R through library glmnet (2.0.10), and the generalized linear model function glm from the R stats library.

To investigate the overall classification performance, we randomly partitioned the data set “RESdep” of retained introns into 90%/10% (i.e., 1268/141) training/test data, and randomly sampled the same amount of training/test data from the control “Ctr” data set. We used the training data to learn a logistic regression model with Lasso variable selection and tested it on the test data. Next, to evaluate how well this model classifies specific subsets of “RESdep” introns and retained introns specific for each mutation, we applied the model trained with “RESdep” vs “Ctr” data having held fixed its parameters to the classification of the 141 control test-introns vs. 141 retained introns subsampled from the following sets: (i) “RESdep_ΔPIR10” introns from the “RESdep” set with ΔPIR > 10 in all three mutants (871 introns); (ii) “NMD”, introns from the “RESdep” set predicted to trigger NMD when retained (574 introns); (iii) “bud13”, introns with ΔPIR>15 upon bud13 mutation at 32 hpf (2,363 introns); (iv) “rbmx2”, introns with ΔPIR>15 upon rbmx2 mutation at 48 hpf (2,186 introns); and (v) “snip1”, introns with ΔPIR>15 upon snip1 mutation at 48 hpf (2,675 introns). We repeated this procedure, including model training and classification of test data, 10,000 times and report average ROC curves (Fig. 6A) and average model coefficients for each feature extracted from the trained models. These averages indicate the direction of the effect (e.g. positively [blue] or negatively [red] associated with retention upon RES mutation; Fig. 6B; Supplemental Fig. 10B and Supplemental Table S3).

To study the potential of each feature to contribute to the discrimination between the “RESdep” and “Ctr” intron sets, we randomly partitioned the dataset of retained introns into 90%/10% (i.e., 1268/141) training/test data, and randomly sampled the same amount of training/test data from Ctr. Using the training data, we learned logistic regression models without Lasso variable selection using only a single feature at a time neglecting all other features. The test data were used to determine the AUC. This experiment was repeated 10,000 times and we report average AUCs for each feature in Fig. 6B and Supplemental Fig. 10B. In addition, the fraction of the null deviance that was reduced by each single-feature model was recorded, and the average reductions of the null deviance for each feature are reported in Supplemental Table S3.

## Acknowledgements

We thank H. Codore and K. Bishop for technical help; C.E. Vejnar for providing UCSC tracks, M. T. Lee for initial intron retention analysis, C. Kontur and D. Burkhardt for manuscript editing and all the members of the Giraldez laboratory for intellectual and technical support; Marc Ekker and Joseph Fetcho for providing zebrafish transfgenic lines, Adife Ercan-Sencicek for cloning human *BUD13* ORF, Sophie Bonnal for advise on the intron retention analyses and interpretation and Juan Valcarcel for critical reading of the manuscript. Programa de Movilidad en Áreas de Investigación priorizadas por la Consejería de Igualdad, Salud y Políticas Sociales de la Junta de Andalucía (M.A.M-M.), NIH grants R21 HD073768, R01 HD074078, GM103789, GM102251, GM101108 and GM081602 (A.J.G.), supported this work. A.J.G is supported by the HHMI Faculty Scholar program, the March of Dimes, the Yale Scholars Program and the Whitman fellowship funds provided by E.E. Just, L.B. Lemann, E. Evelyn and M. Spiegel, the H. Keffer Hartline and E.F. MacNichol Jr at the Marine Biological Laboratory in Woods Hole, MA. M.I. is supported by the European Research Council (ERC) under the European Union's Horizon 2020 research and innovation program (grant agreement No ERC-StG-LS2-637591) and the Spanish Ministry of Economy and Competitiveness (BFU2014-55076-P, and the ‘Centro de Excelencia Severo Ochoa 2013-2017’, SEV-2012-0208 to the CRG).

## Author contributions

J.P.F., M.A.M-M. and A.J.G. conceived the project and J.P.F., M.A.M-M., M.I. and A.J.G. designed the research. J.P.F. and M.A.M-M. performed all experiments and S.H.C. performed confocal imaging. M.I. and A.G carried out computational and statistical analysis. J.P.F., M.A.M-M., M.I. and A.J.G. performed data analysis and wrote the manuscript with input from the other authors. All authors reviewed and approved the manuscript.

---

**Supplemental Figure S1.**
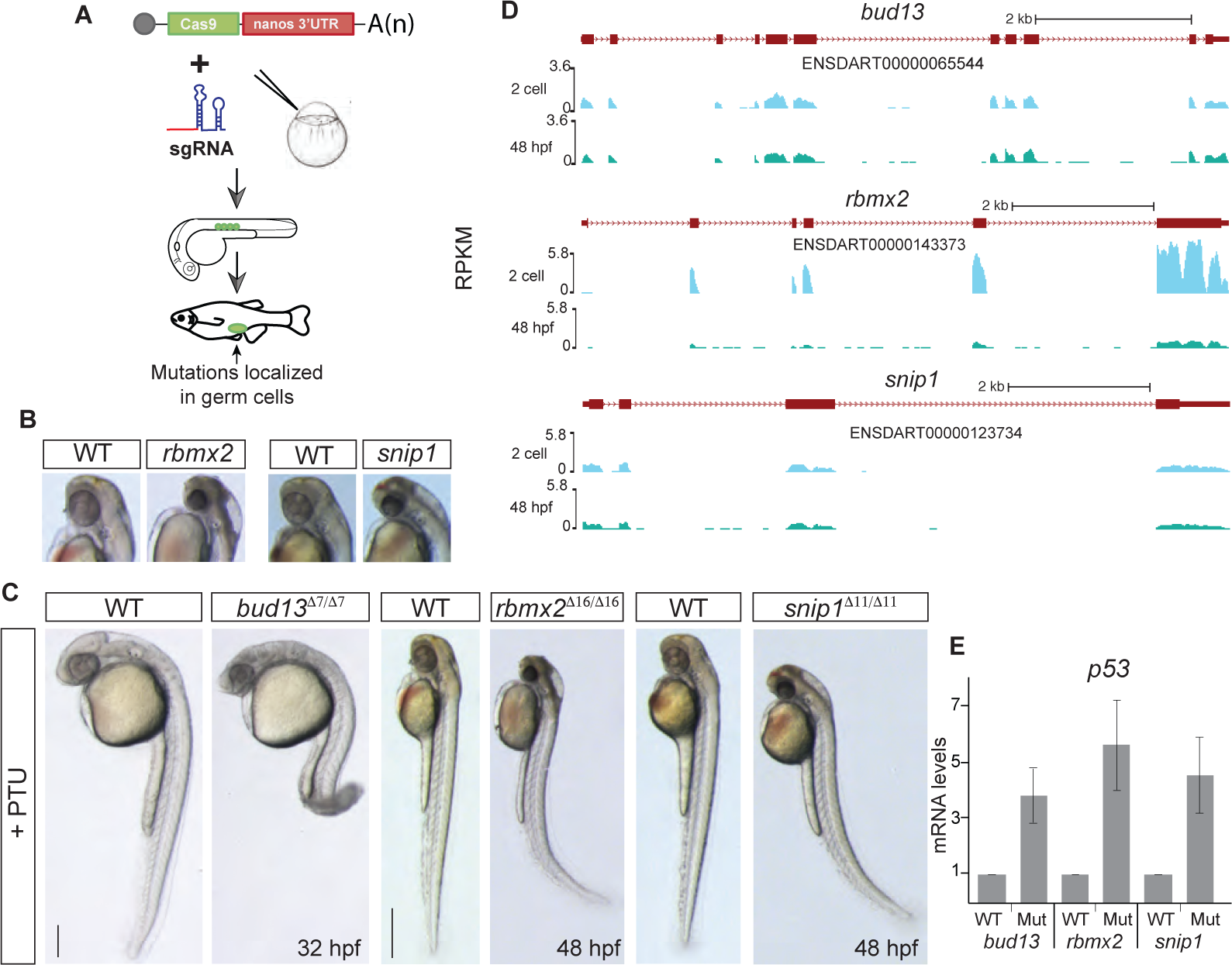
RES complex is required during zebrafish development. A) Scheme illustrating the Cas9-nanos 3′-UTR strategy (Moreno-Mateos et al. 2015). The nanos' 3′-UTR concentrates the expression of Cas9 in the germ cells (green circles). B-C) Bright field microscopy of RES mutant embryos and their corresponding phenotypically wild type sibling (Wt), treated with PTU to avoid melanocyte production, in lateral view (C) or magnification (B, for *rbmx2* and *snip1*). Increased levels of apoptosis, predominantly in the head, are observed upon RES loss-of-function. (scale bar: 1mm at 48 hpf; 0.5mm at 32 hpf). D) UCSC genome tracks showing mRNA levels of RES complex members at 2 cell and 48 hpf stages. E) RT-qPCR showing p53 mRNA levels. Error bars represent SD of the mean from two independent biological replicates (n = 10 embryos per biological replicate). A *p53* up-regulation in the mutants compared to phenotypically WT siblings correlates with the increased cell dead observed in the brain.

**Supplemental Figure S2.**
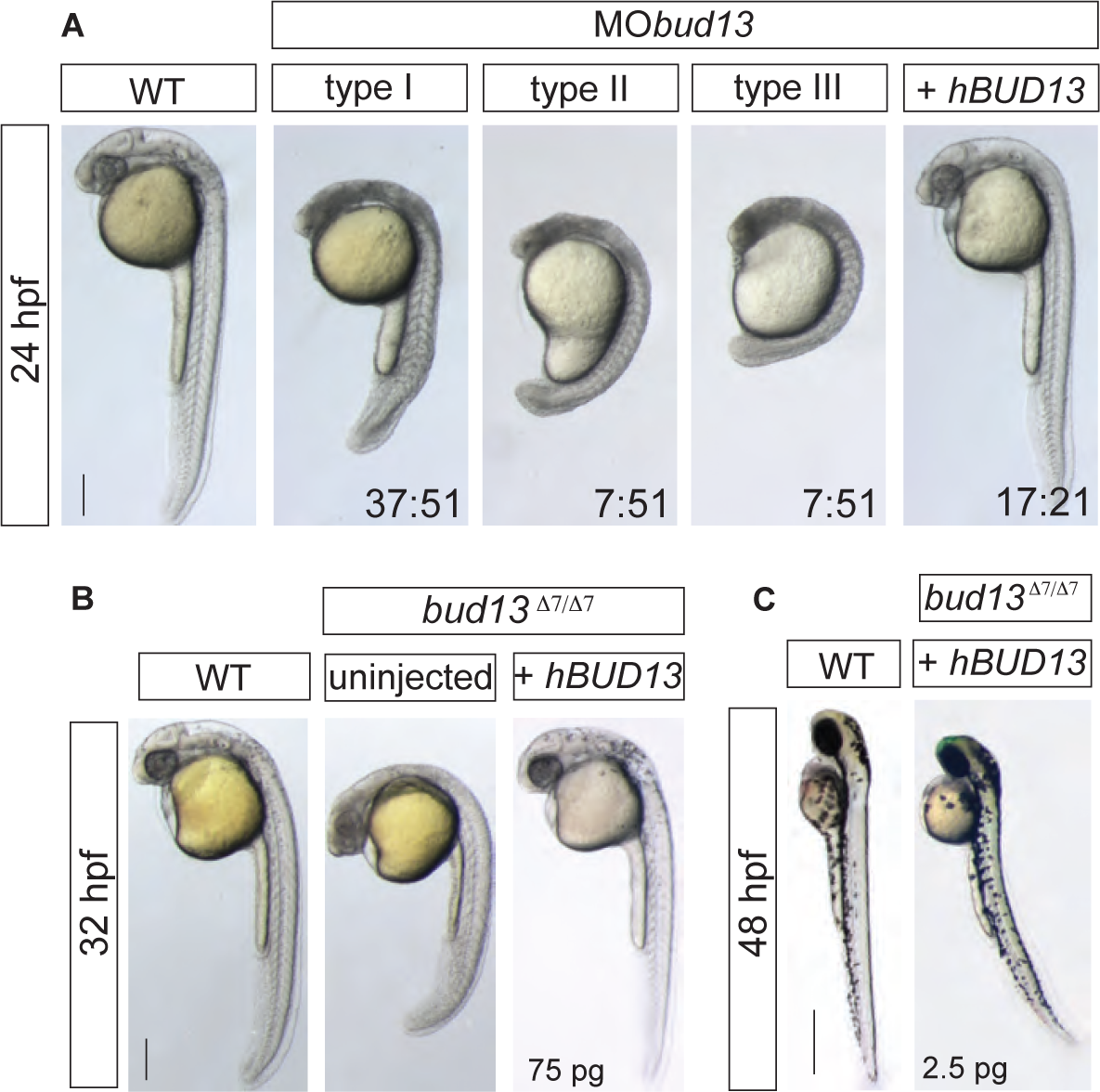
bud13 knock-down show stronger cell death phenotype. A) Lateral view of WT embryos injected with 0.6mM of morpholino antisense oligonucleotide against *bud13* mRNA (MO*bud13*) showing different levels of developmental defects (types I to III). Phenotypes are fully rescue with human *hBUD13* mRNA. (scale bar: 0.5mm). WT: represent phenotypically wild type sibling from the same mutant fish line. Stronger phenotype is likely due to a depletion of the maternal contribution. B) *bud13* mutant embryos fully rescued by providing 75 pg of *hBUD13* mRNA, suggesting that Bud13 function may be conserved across vertebrates (scale bar: 0.5mm). WT: represent phenotypically wild type sibling from the same mutant fish line. C) 48 hpf, *bud13*^Δ7/Δ7^ embryos showing a similar phenotype to *rbmx2*^Δ16/Δ16^ and *snip1*^Δ11/Δ11^ when partly rescued by injection of 2.5 pg of *hBUD13* mRNA (scale bar: 1mm). WT: represent phenotypically wild type sibling from the same mutant fish line.

**Supplemental Figure S3.**
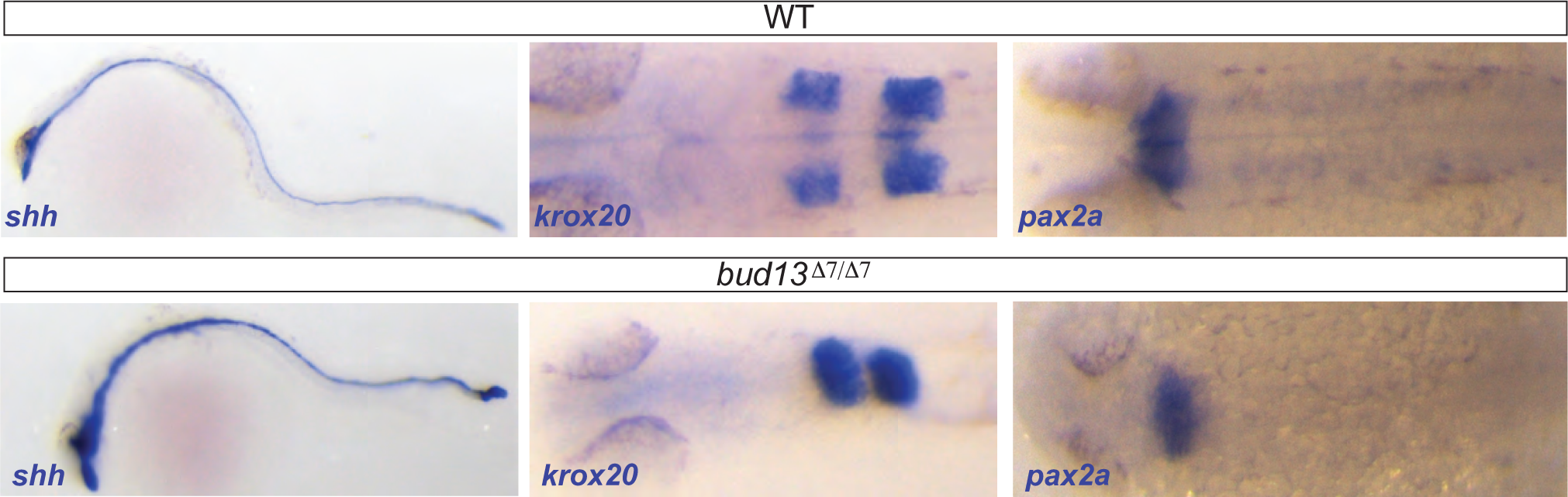
CNS molecular markers show mild differences in *bud13*^Δ7/Δ7^. *In situ* hybridization showing expression pattern of *shh* (notochord and floor plate), *krox20* (*egr2a*; rhombomere 3 and 5) and *pax2a* (anterior midbrain-hindbrain boundary and hindbrain neurons) in WT (top) and mutant (bottom) embryos.

**Supplemental Figure S4.**
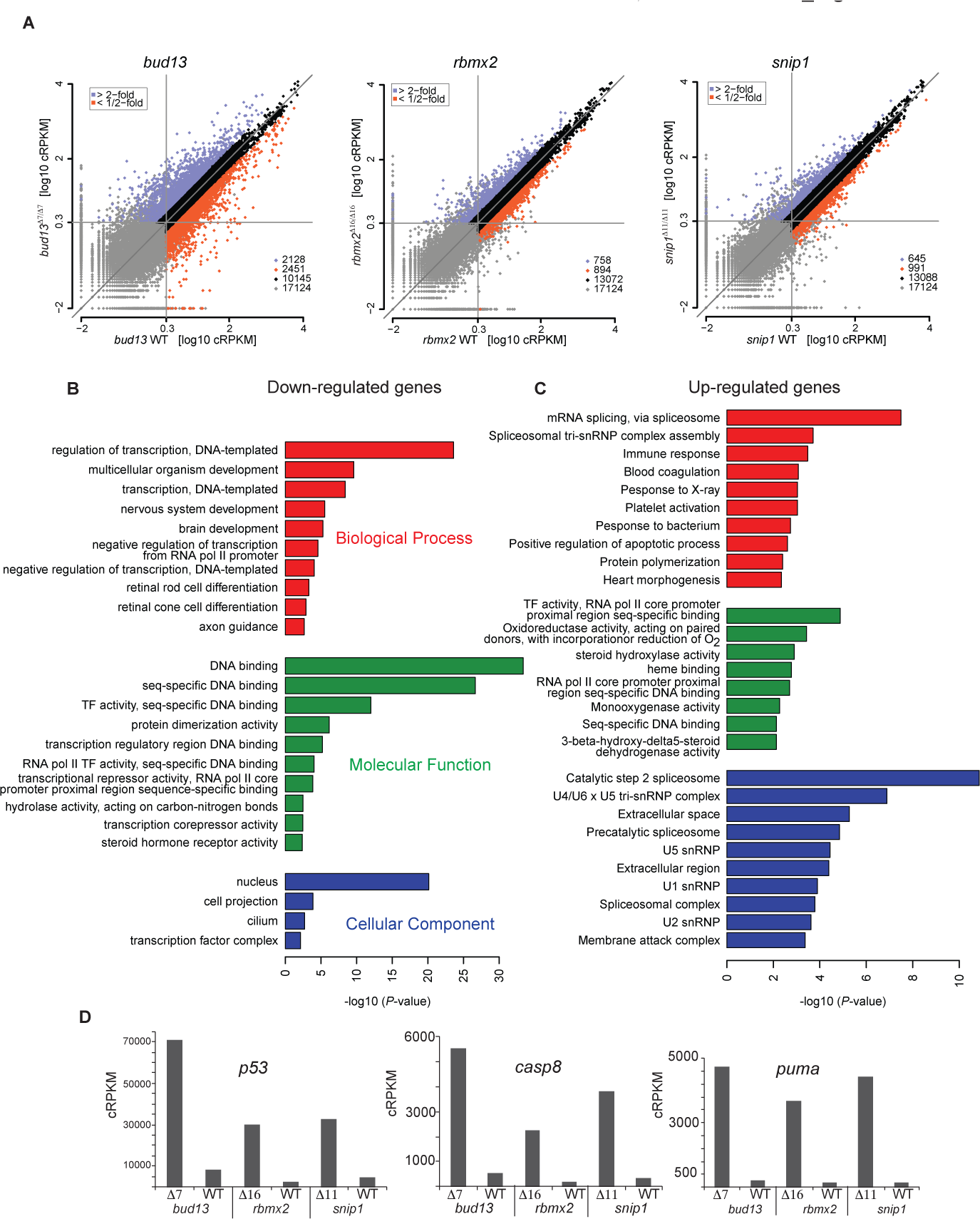
Transcript expression levels and Gene ontology analysis. A) Biplot comparing transcript expression levels in RES mutants and their corresponding phenotypically WT siblings. Genes up-or down-regulation were defined as having a fold change in expression of at least 1.5 in all three mutants and at least 2 for two out the three control vs. mutant individual comparisons (*bud13*, *rbmx2* and *snip1*) (log10 cRPKM). B-C) DAVID cluster analysis of enriched GO annotations for down-regulated (B) or up-regulated genes (C) in RES mutants compared with wild-type siblings. D) Barplots showing expression values (using the cRPKM metric) for up-regulated genes associated with cell death (*p53*, *caspase8* and *puma*). The significant up-regulation in the three mutants correlates with the increased cell dead observed in the developing brain.

**Supplemental Figure S5.**
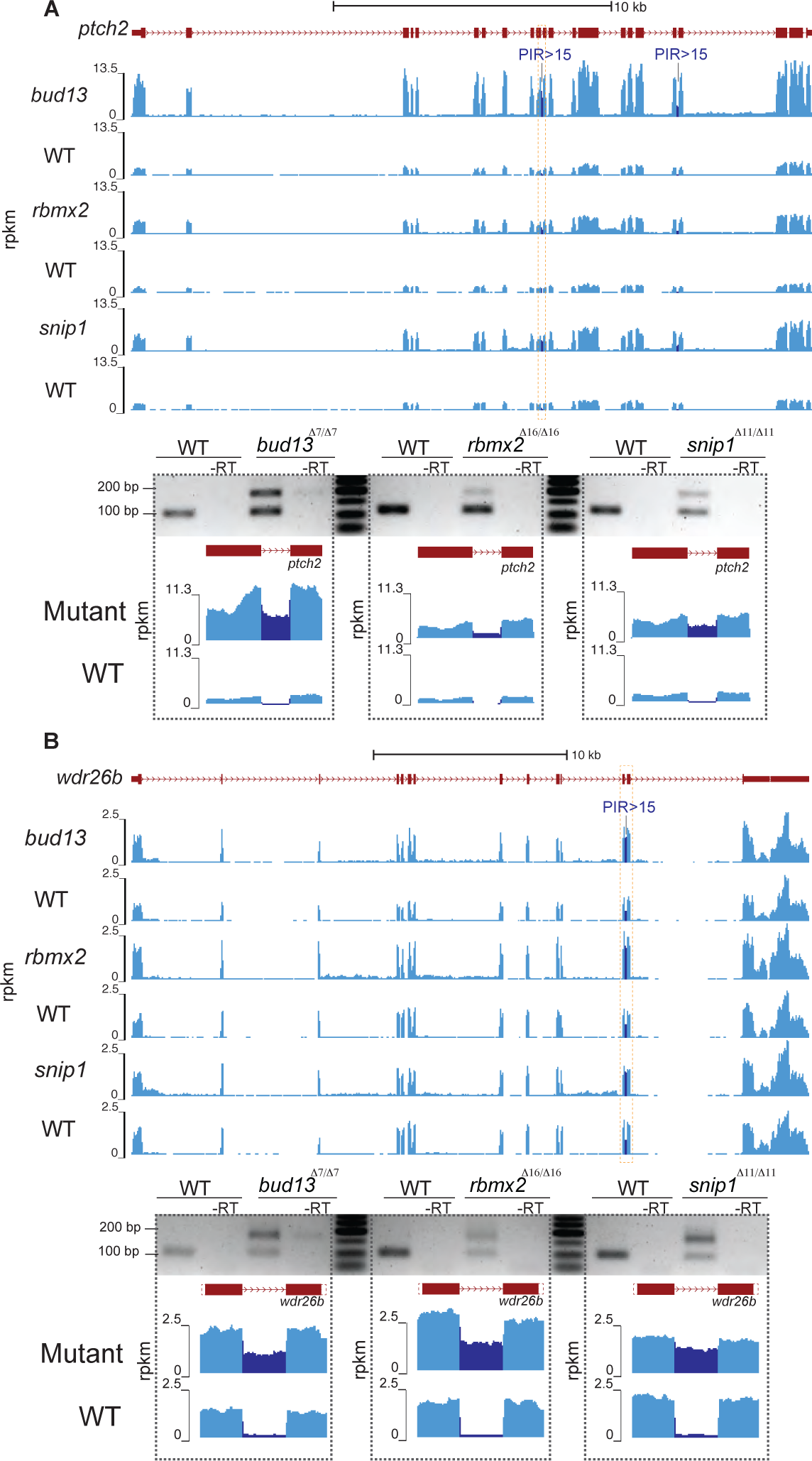
Differentially retained introns detected by RT-PCR. Sequencing read density across the *ptch2* (A) and wdr26b (B) loci (upper panels). RNA-seq signal increases strongly (DPIR>15) in RES mutants only on specific introns (dark blue). RT-PCR assays validating the increased retention (dotted square box) in *bud13*, *rbmx2* and *snip1* mutants compared with the corresponding phenotypically WT siblings (lower panel).

**Supplemental Figure S6.**
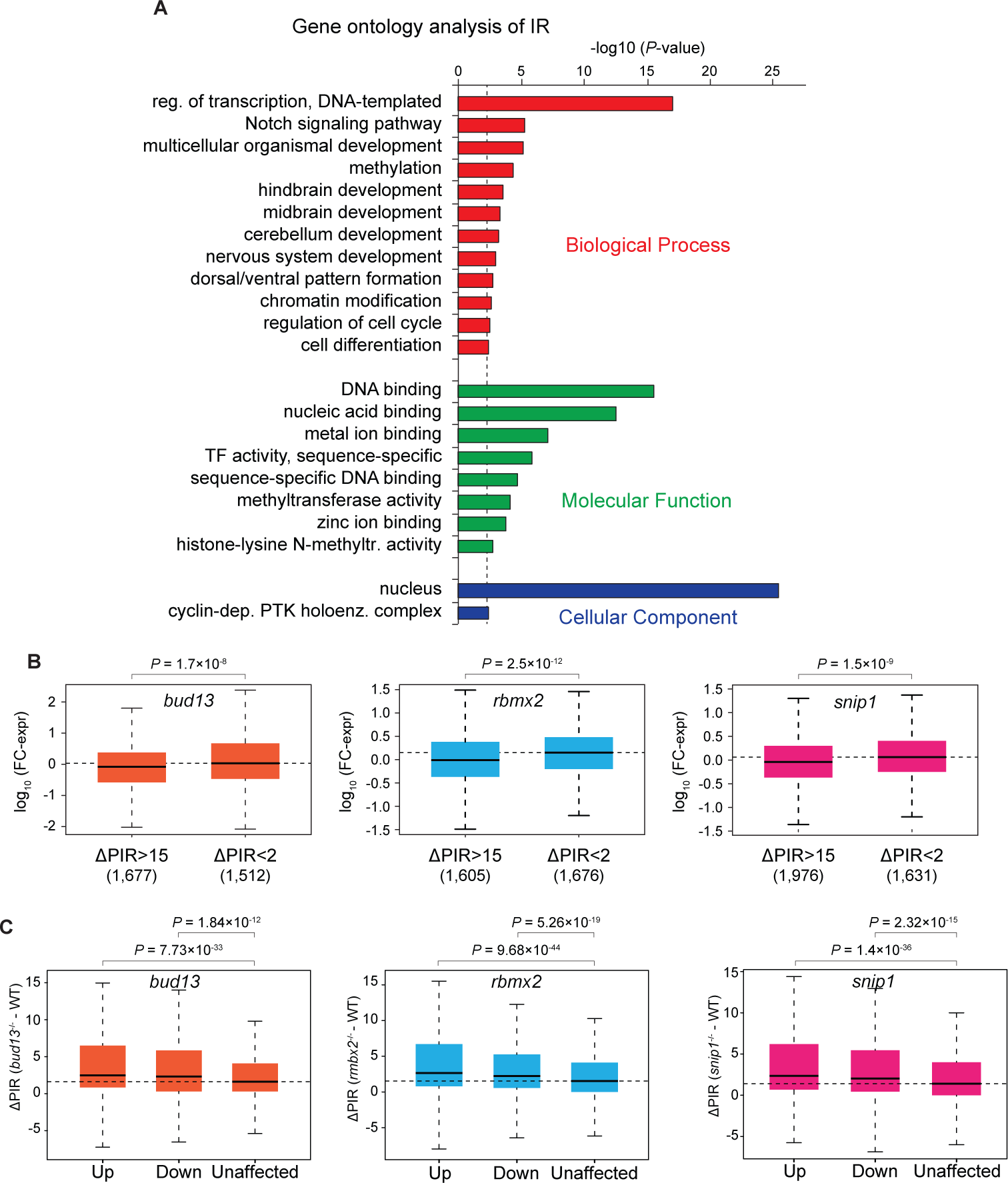
Gene ontology and gene expression analysis of retained introns. A) DAVID cluster analysis of enriched GO annotations for genes that contain introns with highly increased retention (DPIR>15) in at least two of the RES mutants. B) Boxplots showing fold change in expression (FC-expr) for genes containing increased retention (ΔPIR>15) compared with those genes in which all transcripts are not affected (ΔPIR <2) in the RES mutants. *P*-values were calculated using Wilcoxon rank-sum tests. Genes with at least one strongly retained intron (ΔPIR >15) had significantly decreased expression in the mutants compared to genes with no substantial change in intron retention (ΔPIR < 2). C) Boxplots illustrating differences in intron retention between genes that were up-regulated (Up), down-regulated (Down) or did not show significant expression changes (Unaffected). Number of introns in each category: Up = 1,603; Down = 1,623; Unaffected = 69,646. *P*-values were calculated using Wilcoxon rank-sum test. Genes that were differentially expressed in the three mutants (down-or up-regulated) show significantly higher retention (higher ΔPIR) in the mutants compared to their WT-looking siblings.

**Supplemental Figure S7.**
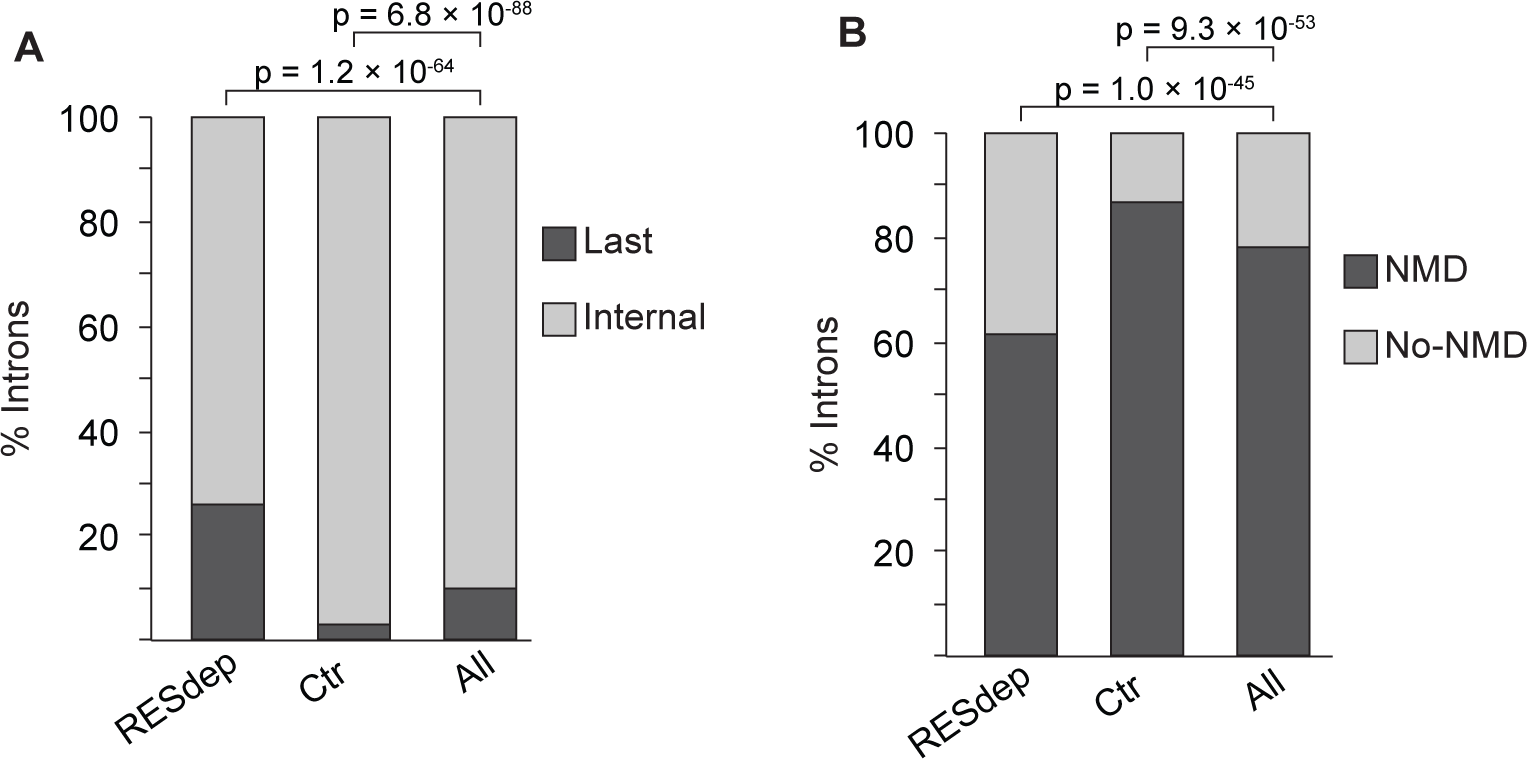
Intron enrichment. Stacked bar plot showing enrichment for last introns (A) and introns predicted not to trigger NMD upon inclusion (B) among the RES-dependent (RES-dep) introns. On the contrary, introns that are not affected by RES depletion (Ctr) are depleted for these types of introns. *P*-values were calculated using Fisher exact test.

**Supplemental Figure S8.**
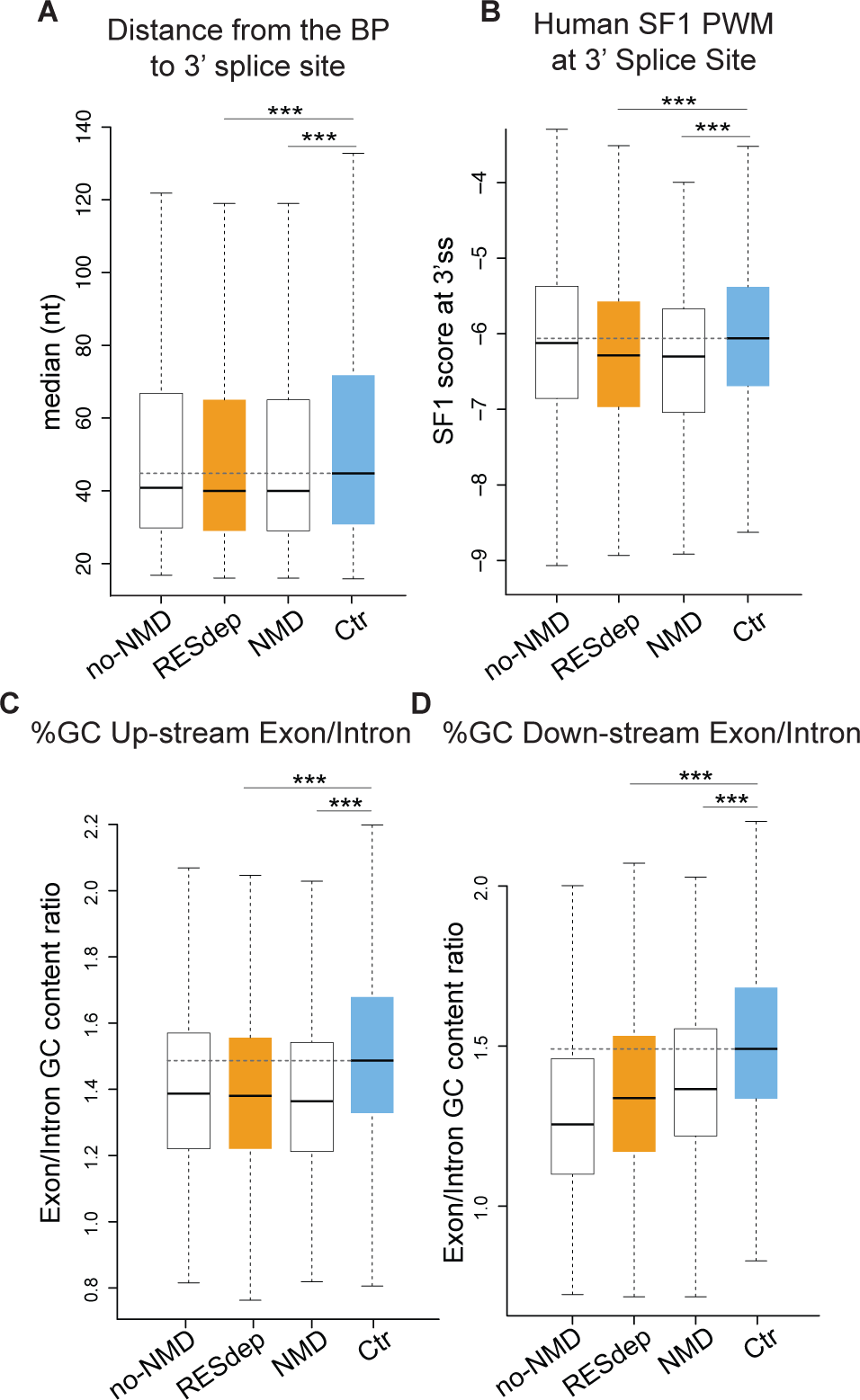
Extended features of the RES-dependent introns. A) Boxplots showing the distribution of the median nucleotide (nt) distance from the top 3 predicted branch points (BP) to the 3' splice site for each intron category. B) Boxplots of the highest score for the human SF1 position weight matrix (PWM) in the 3' intronic region (see Methods for details). C-D) Boxplots showing the GC content ratio between up-stream (C) or down-stream (D) exons vs the retained introns. The lower ratio of no-NMD introns, which are enriched in last introns, is caused by the generally low GC content of last exons overlapping the 3' UTR. (\*\*\**P* ≤ 0.001, Mann-Whitney-U test).

**Supplemental Figure S9.**
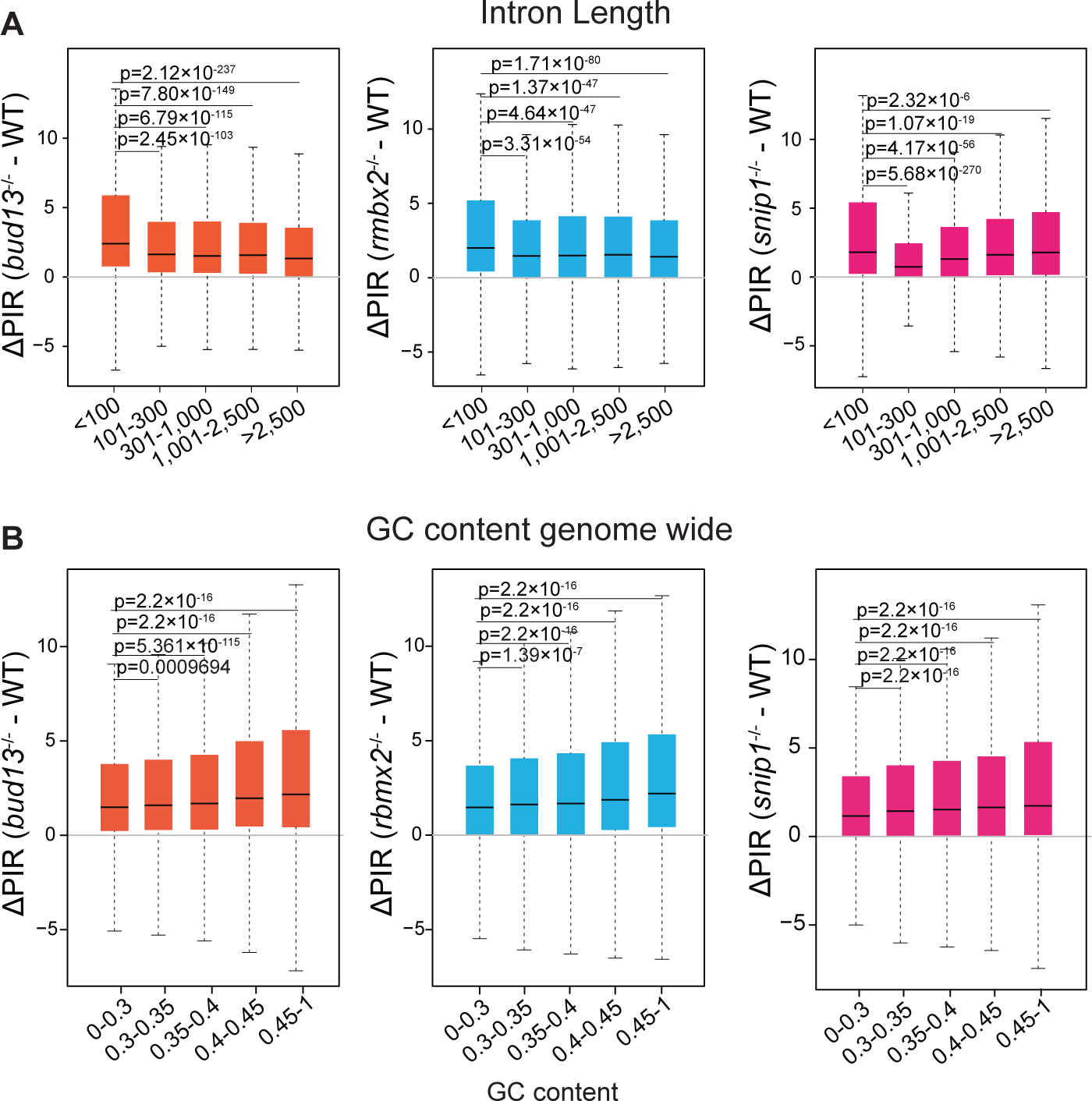
Intron length and GC content genome wide analysis. A) Boxplots showing the degree of change in intron retention (ΔPIR) according to intron length. Number of introns per nucleotide bin: ≤100 = 13,059; 101 – 300 = 13,332; 301-1,000 = 11,808; 1,001 – 2500 = 18,164; >2500 = 16,563. B) Boxplots illustrating the degree of change in intron retention (ΔPIR) according to intron GC content. Number of introns per bin: 0-0.3 = 14663; 0.3-0.35 = 27361; 0.35-0.4 = 21873; 0.4-0.45 =: 6721; 0.45 – 1 = 2308. *P*-values were calculated using Wilcoxon rank-sum test.

**Supplemental Figure S10.**
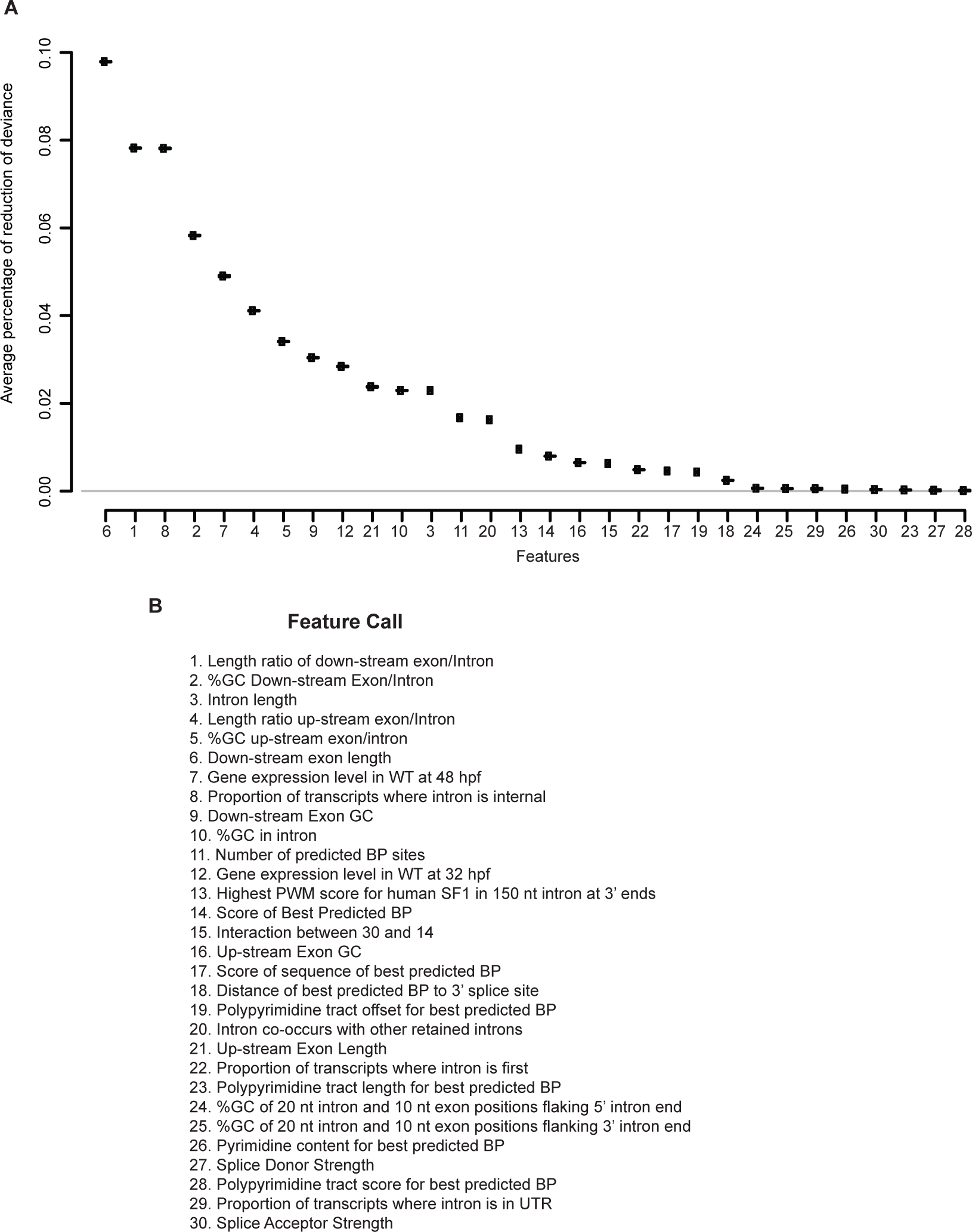
Contribution of each feature to reduction of null deviance. A) Logistic regression models were learned to discriminate between RESdep and Ctr introns with each feature individually and the fraction of the null deviance that was reduced was recorded. Values were averaged over 10,000 repeated holdout experiments. Training data sets consisted of 1,268 RESdep and 1,268 Ctr introns. Error bars indicate 95% confidence interval of reported averages. B) Features call used in Figure 6B and in Figure S10A

